# Modular PEGylation Confers Reduced Immune-Cell Uptake and Enhanced Pharmacokinetic Performance to Encapsulin Protein Nanocages

**DOI:** 10.64898/2026.09.12.751196

**Authors:** Yi Wen Ng, Claire Rennie, India Boyton, Henrico Adrian, Saluo Lai, Daryl Ariawan, Peter Wich, Ole Tietz, Andrew Care

**Affiliations:** School of Life Sciences, University of Technology Sydney, Gadigal Country, Sydney, NSW 2007, Australia; Australian Institute for Microbiology and Infection, University of Technology Sydney, Gadigal Country, Sydney, NSW 2007, Australia; School of Chemical Engineering, University of New South Wales, Sydney, NSW 2052, Australia; Dementia Research Centre, Macquarie Medical School, Macquarie University, NSW 2109, Australia; ARC Centre of Excellence in Synthetic Biology, Macquarie University, NSW 2109, Australia

**Keywords:** encapsulin, protein cage, protein nanoparticle, pharmacokinetics, bio- nano interactions, PEGylation, nanoparticle-based drug delivery, immune evasion

## Abstract

Encapsulins are self-assembling prokaryotic protein nanocages with growing potential as systemic drug delivery systems, but their pharmacokinetic behaviour remains poorly characterised, and rapid immune recognition and clearance may limit delivery to target tissues. Here, we show that modular PEGylation of a SpyCatcher-decorated encapsulin *Alkaliphilus metalliredigens* (Am-S) markedly reduces macrophage uptake and extends systemic circulation. Site-directed surface PEGylation using SpyTagged PEG achieved ∼82% conjugation efficiency, corresponding to an estimated average of ∼49 PEG chains per 60-subunit Am-S nanocage, without compromising nanocage assembly, morphology, or colloidal stability. PEGylated Am-S also retained solubility and protein integrity following freeze–thaw cycling and six months of storage. When interacted with RAW 264.7 macrophages *in vitro*, PEGylation substantially reduced nanocage association and internalisation relative to non-PEGylated nanocages. Following intravenous administration in BALB/c mice, PEGylated nanocages exhibited markedly prolonged circulation, with >50% of the injected dose remaining after 1 h compared with 2.6% for non-PEGylated Am-S, and a circulatory half-life of 1 h 43 min. To our knowledge, this represents the first pharmacokinetic characterisation of an encapsulin nanocage. Together, these findings demonstrate that controlled PEGylation can substantially reduce macrophage interactions and prolong encapsulin circulation *in vivo*.

## Introduction

Encapsulins are prokaryotic protein nanoparticles (PNPs) with growing potential as targeted nanoparticle-based drug delivery systems (NDDSs) [1–3]. They self-assemble from identical subunits into porous icosahedral nanocages of three known structural classes (*T* = 1, 60-mer, 18–24 nm diameter; *T* = 3, 180-mer, 30–32 nm; and *T* = 4, 240-mer, ∼42 nm) [1]. Their hollow architecture provides an internal cavity for loading biotherapeutics (e.g., proteins) or small-molecule drugs, while their repetitive outer surfaces can be modified to display targeting ligands [4–7]. Genetic, chemical, and physicochemical engineering strategies have enabled increasingly sophisticated approaches to cargo loading, targeting cellular delivery, and controlled payload release [1, 8]. However, most studies exploring encapsulin-based NDDSs have demonstrated these functions only *in vitro*, typically through assessments of biocompatibility, target-cell uptake, intracellular payload delivery and activity/cytotoxicity.

Despite extensive *in vitro* development, the behaviour of encapsulin-based NDDSs following systemic administration remains poorly characterised, with only a small number of studies evaluating their performance in animal models [9–11]. These reports demonstrate that intravenous encapsulin delivery, tumour accumulation and therapeutic activity are achievable. However, no studies have characterised encapsulin pharmacokinetics, and the factors governing systemic exposure, clearance, and biodistribution remain poorly understood [1]. Defining and controlling these properties will be essential for the rational development of encapsulins into effective targeted NDDSs suitable for clinical translation.

This translational gap is particularly important because encapsulin-based nanovaccine development is relatively advanced, whereas their use as systemic drug carriers remains less mature due to the distinct biological demands placed on the nanocage [12–15]. Nanovaccines intentionally exploit the inherent immunogenicity of encapsulins to enhance antigen-specific immune responses [9, 16], whereas systemic drug carriers must minimise any immune recognition to remain in circulation long enough to reach distal target tissues (e.g., tumours) [17, 18]. Like other PNPs, encapsulins are susceptible to serum-protein adsorption, opsonisation, and mononuclear phagocyte system (MPS) clearance, which can shorten systemic exposure and limit delivery [9]. They may also elicit particle-specific antibodies that promote immune neutralisation or accelerated clearance upon repeat administration. Consistent with these immune barriers, we previously showed in a preclinical mouse model that intravenously administered *T* = 1 encapsulin derived from *Thermotoga maritima* (TmEnc) was well tolerated, but nevertheless formed a dynamic protein corona, induced nanocage-specific IgM and IgG antibody responses, and was predominantly sequestered by the liver within 6 h [9]. The particles were subsequently internalised and degraded by Kupffer cells, consistent with MPS-mediated clearance.

PEGylation of PNP surfaces has been used to confer antifouling properties that reduce nonspecific protein adsorption and opsonisation, thereby limiting immune recognition and clearance [19]. In some systems, these effects have been shown to prolong nanoparticle circulation, increasing the opportunity for delivery to target tissues while potentially reducing off-target exposure [20–22]. Although methods for PEGylating encapsulins have previously been reported [23], their effects on immune-cell interactions and pharmacokinetics remain unknown.

In this study, we leverage a *T* = 1 encapsulin nanocage previously engineered with the SpyCatcher–SpyTag split-protein coupling system for modular antigen display in nanovaccine applications [12]. We repurpose this platform for controlled PEG conjugation to the nanocage surface and examined how PEGylation affects encapsulin physicochemical properties, uptake by phagocytic immune cells, and systemic pharmacokinetics *in vivo*.

## Results and discussion

### SpyCatcher–SpyTag coupling enables controlled encapsulin PEGylation

PEGylation can reduce immune recognition and clearance of PNPs, but its effects depend on PEG chemistry, molecular weight, architecture and surface density, as well as particle size and morphology (see **Table S1**). Conventional PEGylation commonly targets accessible lysine or cysteine residues on PNP surfaces, although this offers limited control over conjugation site, efficiency, PEG surface density, and distribution [22, 24, 25].

PEGylation of encapsulins has received comparatively limited investigation. Sonotaki et al. previously conjugated amine-reactive PEG to engineered surface lysine residues of the *T* = 1 encapsulin from *Rhodococcus erythropolis* [23]. However, the method required repeated reagent additions over an extended reaction period and generated subunits bearing variable numbers of PEG chains. Moreover, the study principally examined the effects of PEGylation on nanocage structure and disassembly–reassembly, without evaluating encapsulin interactions with immune cells or their *in vivo* behaviour.

To establish a more controlled PEGylation strategy, we repurposed our previously developed SpyCatcher-decorated *Alkaliphilus metalliredigens* encapsulin nanoscaffold, termed Am-S [12]. This engineered nanocage (*T* = 1, 60-mer) comprises subunits bearing a C-terminal SpyCatcher domain, providing up to 60 defined surface conjugation sites. Am-S has previously supported high-density display of SpyTagged antigens for nanovaccine development [12]. Here, we investigated whether the same conjugation sites could be used to attach SpyTag- functionalised PEG and generate a PEG-coated encapsulin nanocage (**Fig. 1a**).

**Figure 1:**
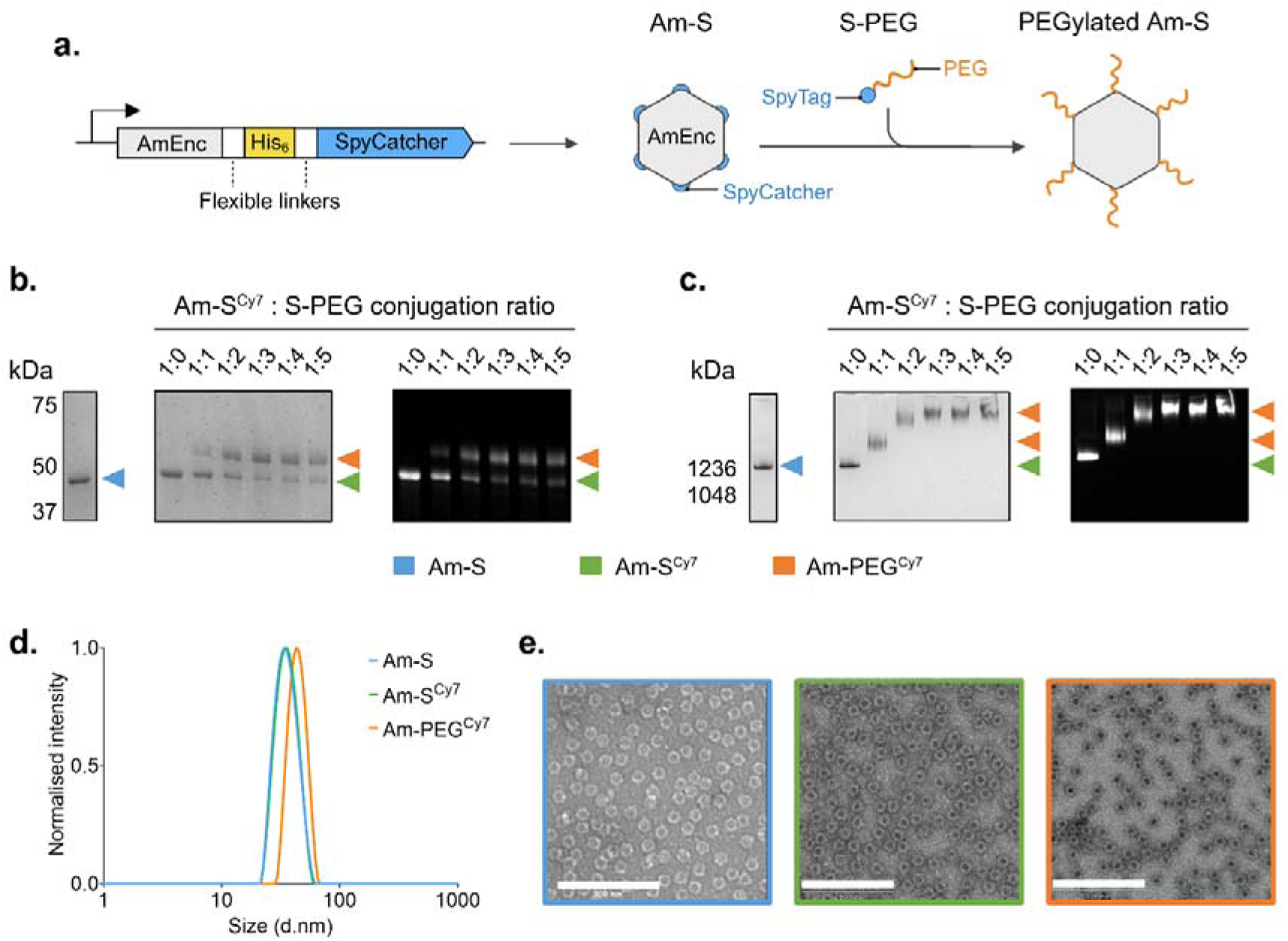
Site-directed PEGylation of the Am-S nanoscaffold through SpyTag– SpyCatcher coupling. **(a)** Schematic of the Am-S genetic construct encoding the *A. metalliredigens* encapsulin subunit (AmEnc) fused at its C-terminus to SpyCatcher via a flexible linker containing a His-tag, and subsequent covalent coupling of SpyTag- functionalised PEG (S-PEG) to self-assembled Am-S nanocages. **(b)** SDS-PAGE analysis of Am-S^Cy7^ conjugation reactions with S-PEG at increasing molar ratios (1:0–1:5): (left) unmodified Am-S subunit reference; (middle) Coomassie staining; and (right) corresponding Cy7 fluorescence. **(c)** BN-PAGE analysis of the same samples: (left) purified unmodified Am-S nanoscaffold; (middle) Coomassie staining; and (right) corresponding Cy7 fluorescence. **(d)** Intensity-weighted DLS size distributions of Am-S (blue), Am-S^Cy7^ (green), and Am-PEG^Cy7^ (orange), showing increased hydrodynamic diameter following PEGylation. **(e)** Negative-stain TEM images of Am-S (blue), Am-S^Cy7^ (green), and Am-PEG^Cy7^ (orange) showing retention of nanocage morphology (Scale bar = 200 nm).

The Am-S nanoscaffold is highly soluble and can be recombinantly produced at yields of ∼200 mg L⁻¹ [12]. Using the previously established workflow, Am-S was expressed in *E. coli* and purified by immobilised metal affinity chromatography with on-column endotoxin removal, followed by size-exclusion chromatography to isolate assembled nanocages (**Fig. S1a–c**). As expected, SDS-PAGE showed a predominant band corresponding to the high purity Am-S subunit (43.9 kDa) (**Fig. 1b**, **left; Fig. S1d**). BN-PAGE showed high-molecular-weight species consistent with nanocage self-assembly (**Fig. 1c**, **left**), while DLS measured a mean hydrodynamic diameter of 34.82 ± 7.47 nm and high monodispersity (PDI = 0.08) (**Fig. 1d**). TEM confirmed a uniform, hollow nanocage morphology matching the Am-S cryo-EM structure (EMD-80547) (**Fig. 1e**) [12].

To enable fluorescence tracking of the PEGylated nanocage in later *in vitro* and *in vivo* experiments, Am-S was NIR dye-labelled with sulfo-Cy7 NHS ester before PEGylation, generating an Am-S^Cy7^ scaffold. Protein-to-dye molar ratios of 1:0.3 to 1:4 were evaluated for degree of labelling and colloidal stability by DLS (**Fig. S2a**). Increasing the dye ratio increased labelling but also produced larger particle populations and higher average PDI values (**Fig. S2b**), consistent with reduced colloidal stability. A 1:0.3 ratio was therefore selected, yielding approximately 12 Cy7 molecules per nanocage while preserving nanocage structure and monodispersity (PDI = 0.08) (**Table S2; Fig. S2b; Fig. 1b-e**).

To enable site-directed PEGylation, SpyTagged PEG (S-PEG) was synthesised by coupling a C-terminal cysteine-bearing SpyTag peptide (2.0 kDa, **Fig. S3a**) to maleimide-functionalised PEG-5000 (∼5.0 kDa) through thiol–maleimide chemistry. MALDI mass spectrometry confirmed S-PEG formation, with an observed average molecular mass of ∼7.4 kDa (**Fig. S3b**). This design enables conjugation of one SpyTagged PEG chain to the single SpyCatcher domain on each Am-S subunit, facilitating a defined 1:1 conjugation stoichiometry. Related SpyTagged PEG reagents have previously been used with SpyCatcher-bearing proteins to construct functional hydrogel systems [26, 27]. However, to our knowledge, their application for nanoparticle PEGylation has not been reported.

Next, to assess the feasibility of SpyTag–SpyCatcher mediated PEGylation and optimise conjugation, Am-S^Cy7^ was reacted with S-PEG at molar ratios of 1:0 to 1:5 for 16 h at 4 °C. Coomassie and red fluorescence imaging of the SDS-PAGE gel showed formation of the PEGylated Am-S^Cy7^ subunit (Am-PEG^Cy7^; ∼51.3 kDa), accompanied by a corresponding decrease in unconjugated Am-S^Cy7^ subunit (43.9 kDa) as the S-PEG ratio increased (**Fig. 1b**). Densitometric analysis determined that subunit conversion reached a maximum of ∼82% at an Am-S^Cy7^:S-PEG ratio of 1:3. BN-PAGE showed a corresponding upward shift of the assembled nanocage band with increasing S-PEG concentration, with an intermediate shift at 1:1 and a plateau from 1:3, consistent with increasing surface PEGylation without nanocage disruption (**Fig. 1c**). The 1:3 ratio was therefore used for all subsequent conjugation reactions.

An ∼82% conjugation efficiency corresponds to an estimated average of ∼49 PEG chains per 60-subunit nanocage. This contrasts with the higher-density but heterogeneous attachment of one to three PEG chains per subunit previously reported for lysine-directed encapsulin PEGylation [23]. Thus, despite incomplete site occupancy, the SpyTag–SpyCatcher system enabled site-specific, 1:1 conjugation of one PEG chain per modified subunit.

Subunit conversion was lower than the relatively complete conjugation efficiency previously achieved using the Am-S platform with smaller SpyTagged peptide antigens, attached either individually or in combination [12]. However, PEGylation of non-Cy7-labelled Am-S showed a similar maximum conversion of ∼73% at a 1:3 ratio (**Fig. S4**), indicating that Cy7 labelling was not the principal cause of incomplete PEGylation of the Am-S^Cy7^ scaffold. The lower conversion achieved with S-PEG may instead reflect steric crowding between polymer conjugates at neighbouring SpyCatcher sites, although this was not directly examined.

PEGylation increased the hydrodynamic diameter of Am-S from ∼34 nm to ∼43 nm and lowered the PDI from 0.05 to 0.02 (**Fig. 1d**), consistent with formation of a hydrated PEG corona. This increase was attributable to PEGylation rather than Cy7 labelling, as Am-S^Cy7^ alone (∼35 nm) was comparable in size to unconjugated Am-S. Negative-stain TEM showed that all three constructs retained the expected nanocage morphology, with comparable protein-core diameters of ∼24 nm (**Fig. 1e**). The smaller TEM diameters reflect visualisation of the dehydrated protein core, whereas DLS measures the solvated particle, including the PEG corona, which is not readily resolved by negative-stain TEM [28].

Collectively, these results establish a facile, site-directed strategy for PEGylating Am-S through SpyTag–SpyCatcher coupling while preserving nanocage assembly and morphology, enabling subsequent evaluation of the effects of PEGylation on encapsulin stability and biological interactions.

### PEGylation supports Am-S stability during freeze–thaw cycling and long-term storage

PNP stability and storability are important attributes for their use as NDDSs, influencing shelf life, handling, and suitability for safe systemic administration. Encapsulin nanocages are inherently stable [29, 30], and we previously showed that unmodified Am-S retained its structure, solubility, and colloidal stability following repeated freeze-thaw cycles and storage for up to six weeks across a broad temperature range [12]. We therefore examined whether Cy7 labelling and subsequent PEGylation altered the stability profile of the Am-S nanoscaffold over an extended timeframe (**Fig. 2**).

**Figure 2.**
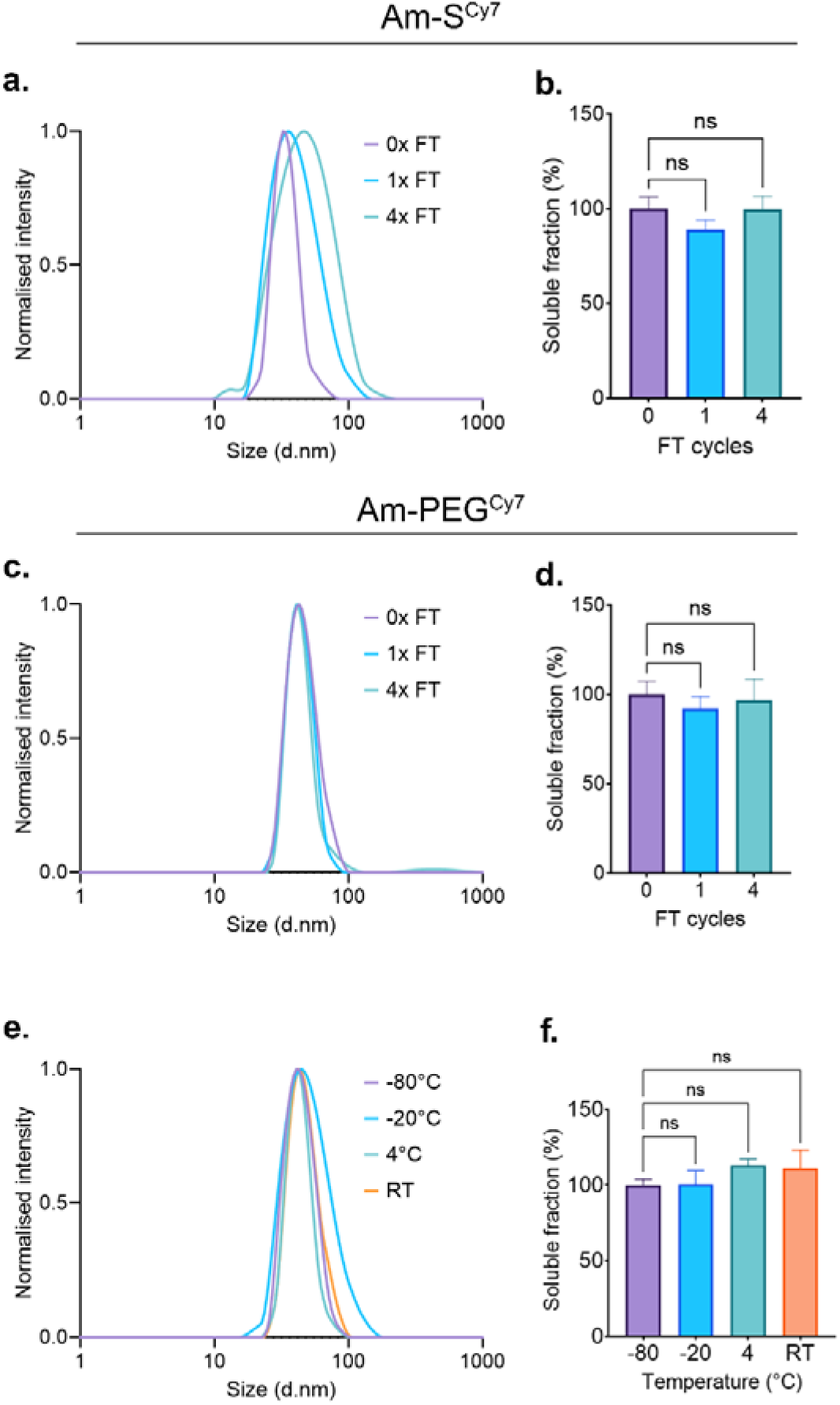
Effect of dye-labelling and PEGylation on Am-S stability and storability. (a-b) Stability of Am-S^Cy7^ following 0×, 1× and 4× freeze–thaw cycles. **(a)** DLS size distributions. **(b)** Soluble protein recovery after freeze–thaw cycling, normalised to untreated Am-S^Cy7^ (0×). **(c-d)** Stability of Am-PEG^Cy7^ following 0×, 1× and 4× freeze–thaw cycles. **(c)** DLS size distributions. **(d)** Soluble protein recovery after freeze–thaw cycling, normalised to untreated Am-PEG^Cy7^ (0×). **(e-f)** Stability of Am-PEG^Cy7^ after six months of storage at −80 °C, −20 °C, 4 °C or room temperature (RT). **(e)** DLS size distributions. **(f)** Soluble protein recovery after storage at the indicated temperatures, normalised to the −80 °C condition. Soluble protein recovery was quantified by SDS-PAGE densitometry. Data are presented as mean ± SD (n = 3). Statistical significance was determined by ordinary one-way ANOVA with Dunnett’s multiple-comparisons test; ns = not significant.

Following one or four freeze–thaw cycles at −80 °C, DLS revealed broader size distributions for Am-S^Cy7^, with the mean hydrodynamic diameter enlarging from 33 nm to 37-42 nm and the PDI increasing from 0.05 to 0.13-0.17 (**Fig. 2a**; **Table S3**). Am-S^Cy7^ nonetheless retained its solubility, with no observable subunit degradation or significant loss of soluble protein (**Fig 2b**; **Fig. S5a**). These changes were not previously observed for unmodified Am-S under comparable conditions, suggesting that Cy7 labelling modestly reduced colloidal stability without compromising protein integrity.

In contrast, Am-PEG^Cy7^ largely retained its colloidal properties following freeze-thaw cycling. Mean hydrodynamic diameters remained between 42 and 44 nm, with low PDI values of 0.06-0.08 across all conditions (**Fig. 2c**; **Table S3**). A low-intensity aggregate population was detected in one sample after four cycles; however, no significant loss of soluble protein or detectable subunit degradation was observed (**Fig. 2d**; **Fig. S5b**). These results indicate that PEGylation mitigated the freeze– thaw-induced changes observed for Am-S^Cy7^.

To assess longer-term stability, Am-PEG^Cy7^ was stored at either room temperature (RT), 4 °C, −20 °C, or −80 °C for six months. DLS revealed temperature-dependent differences in colloidal stability. Samples stored at −20 °C showed greater polydispersity (PDI = 0.16) than those stored at −80 °C, 4 °C, or room temperature (PDI = 0.07–0.1), while one −20 °C sample contained an aggregate population (**Fig. 2e**; **Table S3**). Samples stored at RT also showed a modest increase in mean hydrodynamic diameter to 46 nm, compared with 42 nm at −80 °C and 4 °C. Am- PEG^Cy7^ nevertheless retained its solubility across all temperatures, with no significant differences relative to the −80 °C condition (**Fig. 2f**; **Fig. S5c**). The reduced colloidal stability at −20 °C may reflect freeze-concentration and phase separation during slow freezing [31]. Overall, storage at −80 °C or 4 °C provided the most consistent maintenance of Am-PEG^Cy7^ size distribution and solubility.

### PEGylation reduces macrophage association and uptake of Am-S *in vitro*

PNPs possess repetitive surface architectures and virus-like morphologies that promote uptake by antigen-presenting cells [17, 32]. Although advantageous for vaccination, this can compromise systemic delivery by accelerating immune recognition and clearance [24]. Encapsulin uptake has been reported in macrophage-like cells and murine dendritic cells [5, 33–36], while intravenously administered encapsulins have been shown to be sequestered and processed by hepatic Kupffer cells [9]. Whether PEGylation reduced macrophage interactions with Am-S was therefore investigated (**Fig. 3**).

**Figure 3:**
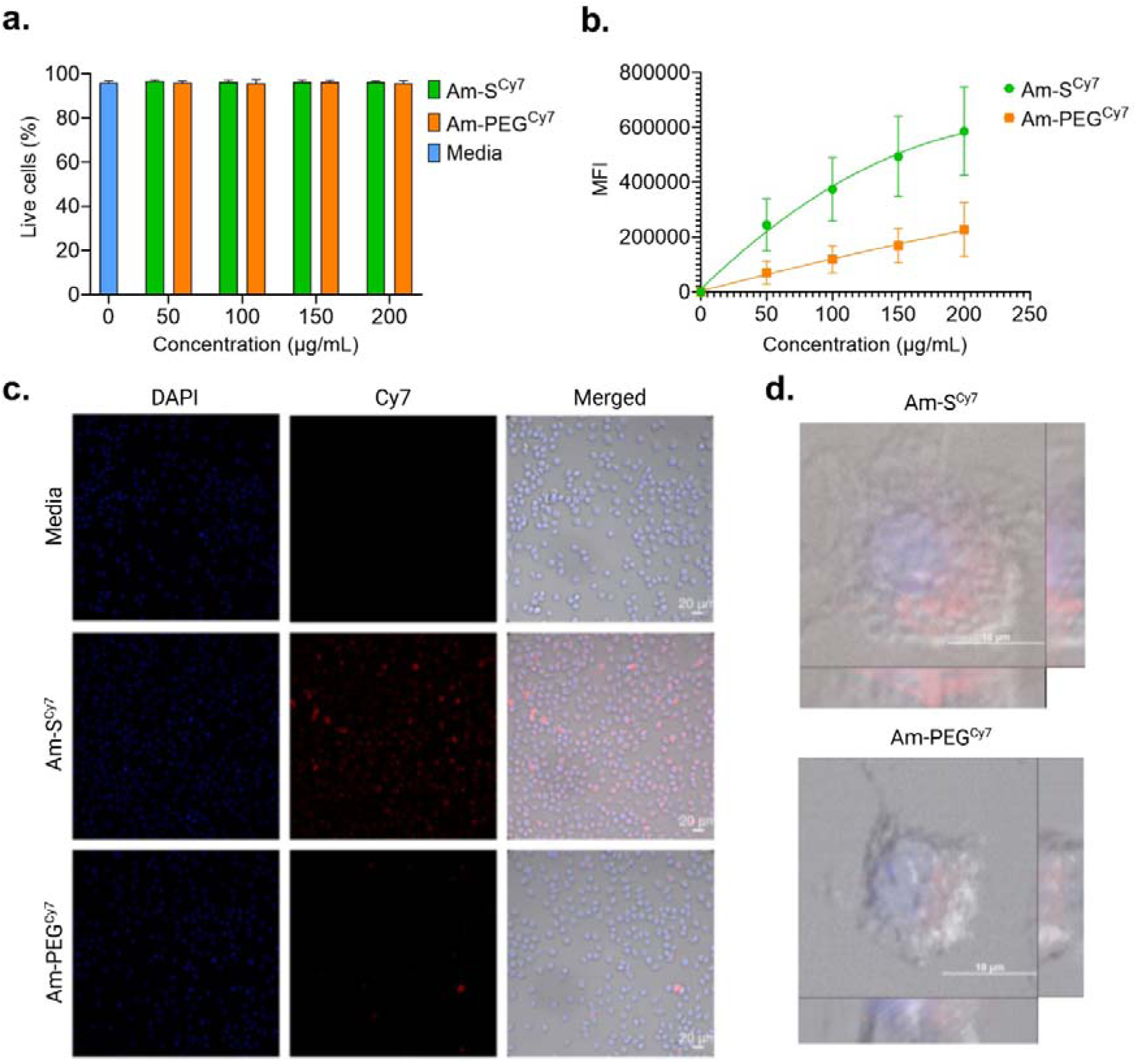
Flow cytometry and microscopy analysis of macrophage association and uptake. **(a)** Viability of RAW 264.7 cells following 2 h exposure to Am-S^Cy7^ or Am-PEG^Cy7^ (0– 200 µg mL⁻¹), relative to media-only controls, as determined by flow cytometry. **(b)** Mean fluorescence intensity (MFI) of cells following 2 h exposure to Am-S^Cy7^ or Am-PEG^Cy7^ across the same concentration range. **(c)** Widefield fluorescence microscopy of cells following 2 h exposure to medium alone (upper), Am-S^Cy7^ (middle), or Am-PEG^Cy7^ (lower). Nuclei stained with DAPI (blue); Cy7 fluorescence shown in red. Scale bars = 20 µm. **(d)** Representative Z- stack images confirming intracellular localisation of Am-S^Cy7^ and Am-PEG^Cy7^ (Scale bars = 10 µm). Data presented as mean ± SD; significance determined by two-way ANOVA with Šídák’s multiple comparisons test; *n* = 3 independent experiments.

Am-PEG^Cy7^ was first incubated in serum-supplemented cell medium at 37 °C for up to 24 h to establish its suitability for cell-based studies. It remained stable throughout, with no evidence of aggregation or fluorophore leaching (**Fig. S6**). Neither Am-S^Cy7^ nor Am-PEG^Cy7^ reduced RAW 264.7 cell viability following 2 h exposure at concentrations up to 200 µg mL⁻¹, with viability comparable to the medium-only control (**Fig. 3a**). This finding is consistent with the favourable *in vitro* biocompatibility previously reported for encapsulin nanocages [4, 7].

Flow cytometry revealed pronounced concentration-dependent association of Am- S^Cy7^ with RAW 264.7 cells (**Fig. 3b**), consistent with previous PNP uptake studies using murine macrophage-like cell lines, including RAW 264.7 and J774A.1 cells [5, 34, 35]. Am-PEG^Cy7^ also showed concentration-dependent association, but at a significantly lower magnitude across all concentrations tested, with the largest differences observed at 150 and 200 µg mL⁻¹ (P < 0.0001). Notably, 200 µg mL⁻¹ Am-PEG^Cy7^ produced a signal comparable to that of only 50 µg mL⁻¹ Am-S^Cy7^. This reduction is consistent with a previous report of an approximately fourfold decrease in RAW 264.7 cell interactions following PEGylation of Tobacco mosaic virus (TMV) PNPs [37].

Because flow cytometry cannot distinguish surface-bound from internalised nanocages, widefield fluorescence microscopy was used to assess cellular uptake after 2 h exposure (**Fig. 3c-d**). Compared to media-only negative controls, Am-S^Cy7^ produced strong, broadly distributed intracellular NIR fluorescence throughout the cell body, whereas substantially less signal was observed following Am-PEG^Cy7^ treatment (**Fig. 3c**). Z-stack imaging demonstrated that the Cy7 signal was localised within cells rather than restricted to the cell surface, verifying nanocage internalisation (**Fig. 3d**).

The robust uptake of Am-S^Cy7^ aligns with previous reports describing efficient phagocytosis of unmodified encapsulins by murine macrophages and dendritic cells, with subsequent trafficking to lysosomes [33]. The diffuse, broadly distributed pattern of Am-S^Cy7^ fluorescence likely reflects lysosomal processing and partial degradation of the nanocage, consistent with this trafficking behaviour. The markedly lower accumulation of Am-PEG^Cy7^ instead indicates that conjugated PEG chains alter the nanocage–cell interface, likely by sterically shielding surface features and limiting the receptor-mediated interactions that drive phagocytic recognition [38]. These *in vitro* studies show that PEGylation reduces macrophage association and internalisation of Am-S.

### PEGylation improves the pharmacokinetic profile of encapsulin nanocages

Having shown that encapsulin PEGylation reduced macrophage association and uptake *in vitro*, its effect on circulatory clearance was next assessed *in viv*o. Healthy BALB/c mice received a single intravenous bolus of Am-S^Cy7^ or Am-PEG^Cy7^ [2.5 mg/kg]. No overt behavioural changes or adverse effects were observed following administration of either formulation, consistent with the tolerability previously reported for subcutaneously administered Am-S [12]. Quantification of the circulating injected dose over time revealed a clear divergence in pharmacokinetic behaviour, with Am-PEG^Cy7^ exhibiting markedly slower clearance than Am-S^Cy7^ (**Fig. 4 and Fig. S7a**).

**Figure 4:**
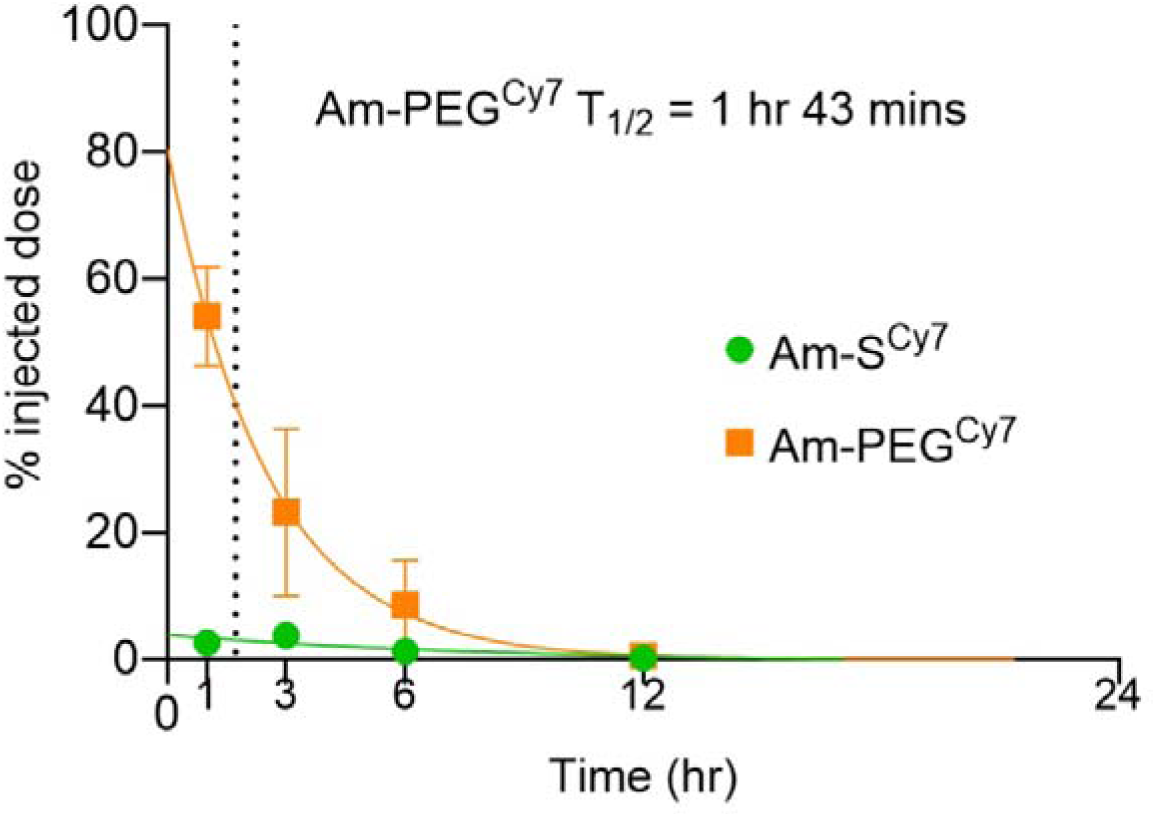
Serum pharmacokinetics of Am-S^Cy7^ and Am-PEG^Cy7^ following IV administration in BALB/c mice. Percentage of injected dose remaining in circulation over time. The T_1/2_ of Am-PEG^Cy7^ was 1 h 43 min; the T_1/2_ of Am-S^Cy7^ could not be reliably determined due to near-background serum fluorescence from the earliest sampled timepoint. Data presented as mean ± SD; *n* = 5–7 mice per timepoint.

The serum concentration–time profile of Am-PEG^Cy7^ is well described by a one-phase decay model, yielding a circulatory half-life (t_1/2_) of approximately 1 h 43 min (**Fig. 4**). By contrast, serum fluorescence associated with Am-S^Cy7^ was at or near background from the earliest sampled time point, precluding reliable model fitting or estimation of a terminal half-life. Am-S^Cy7^ was therefore rapidly cleared from circulation, with only approximately 2.6% of the injected dose remaining after 1 h, compared with more than 50% for Am-PEG^Cy7^. This rapid clearance is consistent with both the pronounced macrophage association and uptake observed for Am-S^Cy7^ *in vitro* and the previously reported MPS-mediated liver sequestration and clearance of systemically administered *T* = 1 TmEnc [9]. Off-target hepatic accumulation of non-PEGylated encapsulins was observed in recent encapsulin-based drug delivery studies with the same TmEnc nanocage [11] and a larger *T* = 3 *Myxococcus xanthus* encapsulin [10], together suggesting that off-target MPS recognition and clearance is a shared feature of non-PEGylated encapsulins *in vivo*. To our knowledge, this work represents the first pharmacokinetic characterisation of an encapsulin nanocage and demonstrates that PEGylation substantially increases its circulatory persistence.

Although PEGylation commonly extends PNP circulation, both native pharmacokinetics and the magnitude of PEG-mediated extension vary substantially, even among particles with similar morphologies (**Table S1**). Many PNPs rapidly accumulate in off-target tissues [39, 40] and exhibit circulatory half-lives of less than 10 min, whereas PEGylation can markedly prolong PNP circulation, with reported increases in half-life ranging from minutes to more than 40 h [21, 25, 41]. Rod-shaped and filamentous PNPs also display variable, frequently biphasic pharmacokinetics irrespective of PEGylation status [20, 41, 42]. These observations indicate that circulatory persistence depends on physicochemical properties beyond particle geometry and PEG conjugation alone.

PEG surface conformation provides a framework linking PEG chain length, grafting density, and biological shielding. Longer PEG chains occupy a larger surface footprint and can reduce the number of chains accommodated per particle; therefore, increasing PEG molecular weight does not necessarily increase shielding if grafting density decreases [43]. PEG chains are commonly classified as adopting mushroom-like or brush-like conformations according to chain spacing relative to the Flory radius [44]. Studies of PEGylated PNPs, including tobacco mosaic virus (TMV), Cowpea mosaic virus (CPMV), and potato virus X (PVX), indicate that PEG conformation can influence antibody recognition, macrophage interactions, and circulation, although these effects remain dependent on the underlying particle system [24, 45, 46].

Based on the experimentally determined conjugation density (∼82%) and known dimensions of Am-S, the attached PEG5000 on the Am-PEG^Cy7^ surface was predicted to adopt a brush-like conformation. This estimate does not account for nanocage surface curvature or heterogeneous PEG occupancy. Brush conformations typically provide enhanced steric shielding and more effectively reduce protein adsorption, immune-cell interactions, and MPS-mediated clearance than mushroom-like arrangements [20, 24, 42, 47]. Am-PEG^Cy7^ exhibited reduced macrophage interactions and substantially prolonged circulation compared to non-PEGylated controls. These findings suggest that the brush-like PEG layer contributed to shielding the encapsulin surface and altering its bio–nano interactions and pharmacokinetics.

The *T* = 1 icosahedral architecture of Am-S provides 60 defined PEG attachment sites and thereby sets the maximum achievable surface density. Conjugation plateaued at approximately 82%, while Cy7 labelling did not measurably reduce PEGylation efficiency. The incomplete occupancy is therefore consistent with steric hindrance, heterogeneous site accessibility, and geometric constraints imposed by nanocage curvature. Despite these constraints, PEGylation of the encapsulin resulted in substantial pharmacokinetic enhancement.

## Conclusions

This study provides the first pharmacokinetic characterisation of an encapsulin protein nanocage and demonstrates that PEGylation can substantially prolong its retention in circulation. Am-S was PEGylated by conjugating SpyTagged PEG to SpyCatcher domains displayed on the nanocage surface, without compromising assembly, morphology or colloidal stability. PEGylated Am-S also showed reduced macrophage association and internalisation *in vitro* and markedly slower clearance following intravenous administration in mice. These findings address a recognised translational limitation of encapsulin-based drug delivery, for which pharmacokinetics and strategies to reduce immune recognition remain poorly defined [1, 9].

The SpyCatcher–SpyTag system provides a controlled approach for engineering Am-S surface composition. Previous studies have shown that this ligation strategy can support the display of functional proteins and targeting ligands on encapsulins [48–50]. Defined conjugation sites could therefore enable co-display of antifouling polymers and targeting ligands while retaining the internal cavity for protein, small-molecule or nucleic-acid loading [1, 4, 5, 34]. PEG and ligand densities could potentially be adjusted to balance circulation time, target binding, and cellular uptake. The same conjugation strategy could also support evaluation of emerging PEG alternatives intended to mitigate anti-PEG antibody-mediated accelerated blood clearance [51].

Because encapsulins comprise a structurally and sequence-diverse family of PNPs, this approach may also be transferable to other scaffolds [52, 53]. We previously observed no antibody cross-reactivity between the *T* = 1 TmEnc and a larger *T* = 4 encapsulin from *Quasibacillus thermotolerans*, supporting potential immune orthogonality [9]. Sequential administration of antigenically distinct encapsulins may therefore offer a complementary strategy for reducing nanocage-specific antibody recognition during repeat dosing, although this remains to be established experimentally.

Further studies should determine how PEG molecular weight, surface density, and conformation affect protein-corona composition, biodistribution, immunogenicity and repeat-dose pharmacology. Cargo-loaded and actively targeted Am-S constructs should also be evaluated in disease-relevant models to determine whether prolonged circulation improves tissue accumulation and therapeutic performance. These findings establish controlled surface PEGylation as a viable strategy for reducing rapid encapsulin clearance and provide a foundation for developing long-circulating, multifunctional encapsulins as targeted NDDSs.

## Materials and methods

### Am-S nanocage production and purification

*E. coli* BL21(DE3) cells (New England Biolabs) carrying pET-24(+) encoding Am-S were cultured overnight in Luria–Bertani (LB) broth containing kanamycin (50 µg/mL) at 37 °C and 200 rpm. Starter cultures were diluted 1:100 (v/v) into 300 mL LB cultures containing kanamycin and grown under the same conditions to an OD_600_ of 0.5–0.6. Am-S expression was induced with 0.1 mM isopropyl β-D-1- thiogalactopyranoside (IPTG), followed by overnight incubation at 20 °C and 200 rpm. Cells were harvested at 10,000 × g for 10 min at 4 °C, and pellets were stored at −30 °C.

Cell pellets were resuspended at 5 mL/g in lysis buffer [20 mM sodium phosphate, 500 mM NaCl, 40 mM imidazole, 25 U/mL Benzonase nuclease, cOmplete Mini EDTA-free Protease Inhibitor Cocktail (Roche; one tablet per 30 mL), and 1 mM phenylmethanesulfonyl fluoride (PMSF), pH 7.4] and incubated on ice for 15 min. Cells were lysed by sonication (Qsonica; 50% amplitude; four cycles of 10 s on/20 s off, with 5 min rest between cycles). Lysates were clarified by centrifugation at 10,000 × g for 15 min at 4 °C and sequentially filtered through 0.45 and 0.2 µm Minisart syringe filters (Sartorius).

Am-S was purified by immobilised metal affinity chromatography (IMAC) using a 5 mL HisTrap High Performance column (Cytiva) on an ÄKTA pure system at 5 mL/min. Bound protein was washed with 50 column volumes of buffer containing 0.1% Triton X-114 for endotoxin reduction, followed by stepwise elution with increasing imidazole concentrations. Am-S eluted at 48% buffer B (240 mM imidazole). Eluted fractions were further purified by size-exclusion chromatography (SEC) using a HiPrep 26/60 Sephacryl S-500 column (Cytiva) at 1 mL/min. Fractions corresponding to assembled Am-S nanocages were pooled and concentrated to 1 mg/mL using 100 kDa Amicon Ultra centrifugal filters (Merck). Protein concentration was determined by Bradford assay, and purified Am-S was stored at −80 °C.

### Synthesis of SpyTag peptides

SpyTag peptide containing a cysteine residue on the C-terminus for downstream PEGylation was synthesized at Macquarie University, Sydney, Australia, using standard Fluorenyl methoxycarbonyl (Fmoc)-solid-phase peptide synthesis on a CEM Liberty Blue™ Peptide Synthesiser (CEM, USA). Briefly, rink amide resin was pre-swelled in 50/50 dimethylformamide (DMF) and dichloromethane (DCM) for 1 hour. Amino acids were dissolved in DMF at a concentration of 0.2 M before being transferred to the synthesiser. Peptide was synthesized using sequential amide coupling from C to N-terminus for 3 minutes at 90°C, using five equivalents of amino acid with 10 equivalents of activator (0.5M DIC (N,N′-Diisopropylcarbodiimide) in DMF) and 5 equivalents of activator base (0.5 M Oxyma (Ethyl cyanohydroxyiminoacetate), 0.05 M DIPEA (N,N-Diisopropylethylamine) in DMF), followed by Fmoc deprotection in 20% piperidine in DMF for 2 minutes at 90°C and resin wash in DMF. Following final Fmoc deprotection, resin was removed from synthesiser, transferred to a syringe fitted with a propylene filter, and washed with DMF, DCM, and methanol before cleavage. Peptide was cleaved from resin using cleavage cocktail of 92.5% TFA (trifluoroacetic acid), 2.5% TIPS (triisopropylsilane), and 2.5% H_2_O for 3 hours at room temperature, precipitated in ice cold diethyl ether, dissolved in H_2_O, freeze dried, and purified using a Shimadzu LC-20AD High-performance liquid chromatography (HPLC, Shimadzu, Japan). Mass spectra were obtained on a Shimadzu LCMS-8050 system (Shimadzu, Japan) in positive electron spray {ESI+} mode, fitted with a Polaris 3 C18-A 150 x 4.6 mm column (Agilent Technologies, USA).

### PEGylation of SpyTag peptides

SpyTag peptide containing a cysteine residue was PEGylated by thiol–maleimide coupling. SpyTag (10 mg) was dissolved in a minimal volume of dimethyl sulfoxide (DMSO), followed by addition of 2 mL phosphate-buffered saline (PBS) and 25.2 mg mPEG5000-maleimide (mPEG5000-Mal; Huateng Pharma) in a molar ratio of 1:1. The reaction was incubated overnight at room temperature with stirring. PEGylated SpyTag (S-PEG) was purified by dialysis using a 3.5 kDa molecular weight cut-off membrane. Matrix-Assisted Laser Desorption/Ionization Time-of-Flight (MALDI-TOF) indicated an average molecular mass of 7.38 kDa. S-PEG was lyophilised and stored at −30 °C until use, then reconstituted in PBS for subsequent conjugation reactions.

### Optimisation of Cy7 dye labelling

Am-S was labelled with sulfo-Cyanine 7-NHS ester (sulfo-Cy7-NHS; Lumiprobe) by NHS ester-mediated amine coupling. Am-S in 100 mM sodium bicarbonate buffer (pH 8.3) was reacted with sulfo-Cy7-NHS at Am-S:dye molar ratios of 1:4, 1:2, 1:1, 1:0.5, and 1:0.3 for 1 h at room temperature with agitation, protected from light. Samples were buffer-exchanged into PBS to remove unreacted dye and centrifuged at 16,900 × g for 30 min at 4 °C to remove precipitates.

The degree of labelling was determined spectrophotometrically using absorbance at 280 nm and the Cy7 absorbance maximum, with correction for dye absorbance at 280 nm (correction factor 0.04; **Table S2**). Labelled nanocages were monitored by DLS following centrifugation to assess colloidal stability. The 1:0.3 Am-S:dye ratio was selected for subsequent PEGylation.

### Optimisation of nanocage PEGylation

Am-S^Cy7^ was reacted with S-PEG at Am-S^Cy7^ subunit:S-PEG molar ratios of 1:1, 1:2, 1:3, 1:4, and 1:5, with the Am-S^Cy7^ concentration held constant. Reactions were incubated overnight at 4 °C with agitation, protected from light, then centrifuged at 16,900 *×* g for 30 min at 4 °C to remove precipitates. The soluble fractions were subjected to four rounds of buffer exchange into PBS using 100 kDa MWCO Amicon Ultra centrifugal filters (Merck) to remove excess unconjugated S-PEG. Samples were analysed by SDS-PAGE and BN-PAGE to assess PEG conjugation and nanocage assembly. The 1:3 ratio was selected for subsequent conjugation reactions.

### Polyacrylamide gel electrophoresis

#### Sodium dodecyl sulfate-polyacrylamide gel electrophoresis (SDS-PAGE)

SDS-PAGE was performed using 10% Mini-PROTEAN TGX precast gels in a Mini-PROTEAN Tetra Cell (Bio-Rad). Samples were mixed 1:1 with 2× Laemmli sample buffer, heated at 99 °C for 10 min, and electrophoresed at 200 V for 35 min in SDS running buffer (25 mM Tris, 192 mM glycine, 1% [w/v] SDS, pH 8.3). Precision Plus Protein Dual Color Standard (Bio-Rad) was used as a molecular weight marker. Cy7 fluorescence was imaged using a ChemiDoc MP Imager (Bio-Rad), after which gels were heat-fixed in water, stained with Coomassie Brilliant Blue G-250 (Sigma), de- stained overnight, and imaged using the Coomassie setting.

#### Blue native PAGE (BN-PAGE)

BN-PAGE was performed using 3–12% Bis-Tris NativePAGE gels in an XCell SureLock Mini-Cell system (Thermo Fisher Scientific). Samples were mixed 1:3 with 4× NativePAGE sample buffer and electrophoresed in 1× anode and cathode buffers at 150 V for 1.5 h, followed by 250 V for 1 h. Cy7 fluorescence was imaged using the ChemiDoc MP Imager, after which gels were fixed in 40% methanol/10% acetic acid, de-stained in 10% acetic acid, and imaged using the Coomassie setting.

### Protein quantification

Gel band intensities were quantified by densitometry using ImageJ. Protein concentrations were determined by Bradford assay using BSA standards prepared according to the manufacturer’s instructions (Thermo Fisher Scientific). Standards, blanks, and purified encapsulin samples (5 µL) were analysed in technical triplicate in 96-well plates (Corning) with 250 µL Coomassie Protein Assay Reagent (Thermo Fisher Scientific). Plates were incubated for 10 min with gentle agitation, and absorbance was measured at 595 nm using a Tecan Infinite M200 Pro plate reader. Protein concentrations were calculated from the standard curve.

### Stability of encapsulin nanocages

Freeze–thaw stability was assessed using 70 µL aliquots of Am-S^Cy7^ and Am-PEG^Cy7^ (0.5 mg/mL). Samples were frozen at −80 °C for 15 min and thawed in water at ambient temperature (20-25 °C) for 30 s per cycle, for a total of one or four freeze– thaw cycles. Samples were centrifuged at 16,900 × *g* for 30 min at 4 °C to remove insoluble material, and soluble fractions were analysed by DLS and SDS-PAGE. Unfrozen samples were used as controls and defined as 100% soluble.

Long-term storage stability was assessed using triplicate 100 µL aliquots of Am- PEG^Cy7^ (1 mg/mL) stored at 25, 4, −20, or −80 °C for 6 months. Samples were centrifuged at 16,900 × *g* for 30 min at 4 °C to remove insoluble material. Soluble fractions were analysed by SDS-PAGE to assess protein integrity and soluble recovery, with samples stored at −80 °C defined as 100% soluble. DLS was performed on soluble samples adjusted to 0.15 mg/mL to assess hydrodynamic size and colloidal stability.

Stability in cell culture medium was assessed using a protocol adapted from Diaz et al [4]. Am-S^Cy7^ and Am-PEG^Cy7^ were diluted to 1.9 µM in Dulbecco’s modified Eagle’s medium (DMEM; Gibco) supplemented with 10% fetal bovine serum (FBS) and 1% penicillin–streptomycin and incubated at 37 °C with shaking at 400 rpm for 2, 4, 8, or 24 h. Samples were analysed by SDS-PAGE to assess protein integrity.

### Dynamic Light Scattering (DLS)

Hydrodynamic diameter and polydispersity index (PDI) of encapsulin nanocages were measured before and after PEG conjugation. Samples were centrifuged at 16,900 × g for 30 min at 4 °C to remove insoluble material. Soluble samples (70 µL, 0.15 mg/mL) were loaded into ZEN0040 cuvettes and analysed at 25 °C using a Zetasizer Nano LS (Malvern).

### Transmission Electron Microscopy (TEM)

To visualise the Am-S^Cy7^ and Am-PEG^Cy7^ encapsulin structures, TEM was performed using a TFS Talos 120C microscope operating at 120 kV. For Am-S encapsulin, FEI Tecnai G2 20 microscope was used, operating at 200 kV. 10 μL of sample (200 μg/mL) was placed onto carbon film-coated 300-mesh copper grids (ProSciTech) and subjected to negative staining with a 1:4 dilution of uranyl acetate replacement stain (UAR-EMS) for 1 hour, followed by washing with ultrapure water.

### Quantification of nanocage association with macrophages

RAW 264.7 murine macrophages were cultured in DMEM supplemented with 10% FBS and 1% penicillin–streptomycin at 37 °C with 5% CO₂. Cells were seeded overnight in 24-well plates at 2.5×10 cells/well, followed by overnight serum starvation. Am-S^Cy7^ and Am-PEG^Cy7^ were added in serum-supplemented medium at 50, 100, 150, or 200 µg/mL, with medium-only wells serving as negative controls. Following 2 h incubation at 37 °C with 5% CO₂, cells were harvested, stained with DAPI (5 µL of 10 µg/mL), and analysed using a CytoFLEX S flow cytometer (Beckman Coulter).

Cells were gated by SSC-A/FSC-A, followed by singlet selection (FSC-H/FSC-A) and exclusion of DAPI-positive cells. DAPI-negative, Cy7-positive cells (Cy7 fluorescence >10 ) were classified as associated with Am-S^Cy7^ or Am-PEG^Cy7^. Three independent experiments were performed, with treatments analysed in triplicate.

### Visualisation of nanocage uptake by macrophages

RAW 264.7 macrophages were maintained as described above. Cells were seeded overnight in black 96-well plates at 1×10^5^ cells/well, followed by overnight serum starvation. Am-S^Cy7^ and Am-PEG^Cy7^ (200 µg/mL) were added in serum-supplemented medium and incubated for 2 h at 37 °C with 5% CO₂. Cells were washed three times with PBS, fixed with 4% paraformaldehyde for 15 min at room temperature, washed three times with PBS, and stained with DAPI (10 µg/mL). Following a final PBS wash, cells were imaged using a Nikon ECLIPSE Ti2-E inverted fluorescence microscope. Images were processed using NIS-Elements software (Nikon).

### *In vivo* pharmacokinetic study

Eight-week-old male BALB/c mice (20-26 g) were housed under a 12-hour light/dark cycle with food and water available *ad libitum*. To assess the *in vivo* pharmacokinetics of encapsulins, mice received a single intravenous injection of either of Am-S^Cy7^ or Am-PEG^Cy7^ (2.5 mg/kg) in PBS. Untreated (naïve) control mice did not receive any injection. Non-terminal blood collections were performed *via* submandibular bleeds at early timepoints (1, 3, and/or 6-hour post-injection), with a maximum of two submandibular bleeds per mouse (**Table S4**). Terminal blood collection was performed once per mouse by cardiac puncture under anaesthesia, followed by euthanasia at the predefined timepoints (12 or 24-hour). Untreated control mice did not undergo submandibular bleeds and were subjected to a terminal blood collection only.

### Serum fluorescence analysis

Whole blood collected at each time-point was allowed to clot for 15 min at room temperature and centrifuged at 2,000 × *g* for 10 min at 4 °C. Serum fluorescence was measured in black 384-well plates using a Tecan Infinite M200 Pro plate reader (Excitation: 747 nm; Emission: 780 nm). A serum-based calibration curve was generated by spiking naïve serum with known concentrations of Am-S^Cy7^ (**Fig. S7a**). The same calibration curve was used for Am-S^Cy7^ and Am-PEG^Cy7^, as both showed comparable linear fluorescence responses in naïve serum under identical measurement conditions (**Fig. S7b**). Serum fluorescence values from treated mice were interpolated from the calibration curve and expressed as a percentage of the injected dose.

### PEG conformation calculations

Using the cryo-EM diameter of Am-S (21.2 nm) as an approximation of the grafting surface, the surface area available for PEG attachment was calculated to be 1412 nm^2^ assuming a spherical geometry. Based on an 82% PEGylation efficiency, approximately 49 PEG chains were estimated to be present per nanocage, corresponding to an average inter-chain spacing (D) of 5.37 nm. The Flory Radius (R_F_) of PEG5000 was calculated using the relationship R_F_ = aN^3/5^, where a is the monomer length (3.5 Å) and N is the number of ethylene glycol repeat units (114), yielding an R_F_ of 6.0 nm. PEG conformation was then assessed by comparison of D and R_F_, where D < R_F_ indicates a brush-like conformation and D > R_F_ indicates a mushroom-like conformation.

### Ethical statement

All animal models used were approved by the University of Technology Sydney’s Animal Care and Ethics Committee (ETH22-6924) and were in accordance with the Australian Code for the Care and Use of Animals for Scientific Purposes, 8^th^ Edition, 2013 guidelines.

### Statistical analysis

Statistical significance was determined with Ordinary One-way ANOVA and Two-Way ANOVA using GraphPad PRISM. Results are presented as mean ± standard deviation (SD).

## Acknowledgements

This work was supported by the National Health & Medical Research Council (NHMRC, 2037822), Dementia Australia Research Foundation (DARF), the Mason Foundation, and the National Foundation for Medical Research & Innovation (NFMRI). The authors acknowledge the use of the equipment Nikon ECLIPSE Ti2-E inverted fluorescence microscope in the Microbial Imaging Facility at AIMI in the Faculty of Science, UTS. We would like to thank Amy Bottomley for their scientific input and/or technical assistance. We also acknowledge the use of equipment and infrastructure within the Ernst Facility in the UTS Faculty of Science. We further acknowledge the facilities and the scientific and technical assistance of Microscopy Australia at the Electron Microscope Unit (EMU) within the Mark Wainwright Analytical Centre (MWAC) at UNSW Sydney.

We acknowledge and pay respect to the Gadigal people, the traditional custodians of the land on which this research was conducted.

## References

1. Kwon, S. and T.W. Giessen, Engineering encapsulin nanocages for drug delivery. Materials Advances, 2025. 6(18): p. 6209–6220.

2. Chmelyuk, N.S., et al., Encapsulins: structure, properties, and biotechnological applications. Biochemistry (Moscow), 2023. 88(1): p. 35–49.

3. Rodriguez, J., et al., Nanotechnological applications based on bacterial encapsulins. Nanomaterials 11 (6): 1467. 2021.

4. Diaz, D., et al., Bioengineering a light-responsive encapsulin nanoreactor: a potential tool for in vitro photodynamic therapy. ACS applied materials & interfaces, 2021. 13(7): p. 7977–7986.

5. Coffeen, C.F., et al., Encapsulin-Protected Immunotherapeutic Complexes: Bacteria-Derived Nanoparticles for mRNA Delivery to Eukaryotic Cells. ChemMedChem, 2026. 21(11): p. e202501050.

6. Van de Steen, A., et al., Bioengineering bacterial encapsulin nanocompartments as targeted drug delivery system. Synthetic and Systems Biotechnology, 2021. 6(3): p. 231–241.

7. Moon, H., et al., Developing genetically engineered encapsulin protein cage nanoparticles as a targeted delivery nanoplatform. Biomacromolecules, 2014. 15(10): p. 3794–3801.

8. Jones, J.A., R. Benisch, and T.W. Giessen, Encapsulin cargo loading: progress and potential. Journal of Materials Chemistry B, 2023. 11(20): p. 4377–4388.

9. Rennie, C., et al., In vivo behavior of systemically administered encapsulin protein nanocages and implications for their use in targeted drug delivery. Advanced Therapeutics, 2024. 7(2): p. 2300360.

10. Zhang, Y., et al., Genetically engineered magnetic nanocages for cancer magneto-catalytic theranostics. Nature Communications, 2020. 11(1): p. 5421.

11. Obozina, A.S., et al., Genetically Encoded In Vivo Ligation-Driven Targeted Drug Delivery System for Oncotheranostics. Advanced Healthcare Materials, 2026. 15(9): p. e04119.

12. Boyton, I., et al., Engineering a Novel Bacterial Encapsulin for Programmable Surface Functionalization: From Single-Target to Mosaic Nanovaccines. Biorxiv: the Preprint Server for Biology, 2026.

13. Gorman, J., et al., Cleavage-intermediate Lassa virus trimer elicits neutralizing responses, identifies neutralizing nanobodies, and reveals an apex-situated site-of-vulnerability. Nature Communications, 2024. 15(1): p. 285.

14. Jung, H.-G., et al., Molecular design of encapsulin protein nanoparticles to display rotavirus antigens for enhancing immunogenicity. Vaccines, 2024. 12(9): p. 1020.

15. Wang, Z., et al., Extraordinary titer and broad anti-SARS-CoV-2 neutralization induced by stabilized RBD nanoparticles from strain BA. 5. Vaccines, 2023. 12(1): p. 37.

16. Bhattacharya, S., et al., Heterologous Prime-Boost with Immunologically Orthogonal Protein Nanoparticles for Peptide Immunofocusing. ACS Nano 2024, 18, 20083–20100.

17. Sandra, F., et al., Developing protein-based nanoparticles as versatile delivery systems for cancer therapy and imaging. Nanomaterials, 2019. 9(9): p. 1329.

18. Chung, Y.H., H. Cai, and N.F. Steinmetz, Viral nanoparticles for drug delivery, imaging, immunotherapy, and theranostic applications. Advanced drug delivery reviews, 2020. 156: p. 214–235.

19. The benefits and risks of PEGylation in nanomedicine. Nature Nanotechnology, 2025. 20(5): p. 575–575.

20. Lee, K.L., et al., Stealth filaments: Polymer chain length and conformation affect the in vivo fate of PEGylated potato virus X. Acta biomaterialia, 2015. 19: p. 166–179.

21. Li, C., et al., Real-time monitoring surface chemistry-dependent in vivo behaviors of protein nanocages via encapsulating an NIR-II Ag2S quantum Acs Nano, 2015. 9(12): p. 12255–12263.

22. Huang, X., et al., Hypoxia-tropic protein nanocages for modulation of tumor-and chemotherapy-associated hypoxia. Acs nano, 2019. 13(1): p. 236–247.

23. Sonotaki, S., et al., Successful PEGylation of hollow encapsulin nanoparticles from Rhodococcus erythropolis N771 without affecting their disassembly and reassembly properties. Biomaterials science, 2017. 5(6): p. 1082–1089.

24. Steinmetz, N.F. and M. Manchester, Pegylated viral nanoparticles (VNPs) for biomedicine: the impact of PEG chain length on VNP cell interactions in vitro and ex vivo. Biomacromolecules, 2009. 10(4): p. 784.

25. Hu, H., et al., Physalis mottle virus-like nanoparticles for targeted cancer imaging. ACS applied materials & interfaces, 2019. 11(20): p. 18213–18223.

26. Hammer, J.A., et al., Cell-compatible, site-specific covalent modification of hydrogel scaffolds enables user-defined control over cell–material interactions. Biomacromolecules, 2019. 20(7): p. 2486–2493.

27. Cui, J., et al., Engineering a Thermally Activatable SpyCatcher/SpyTag Protein Ligation for Injectable and In Situ-Forming Hydrogels. Angewandte Chemie International Edition, 2026. 65(3): p. e09477.

28. Harris, J., et al., Application of the negative staining technique to both aqueous and organic solvent solutions of polymer particles. Micron, 1999. 30(4): p. 289–298.

29. Boyton, I., et al., Characterizing the dynamic disassembly/reassembly mechanisms of encapsulin protein nanocages. ACS omega, 2022. 7(1): p. 823–836.

30. Van de Steen, A., et al., Encapsulation of transketolase into in vitro-assembled protein nanocompartments improves thermal stability. ACS Applied Bio Materials, 2024. 7(6): p. 3660.

31. Ball, R.L., P. Bajaj, and K.A. Whitehead, Achieving long-term stability of lipid nanoparticles: examining the effect of pH, temperature, and lyophilization. International journal of nanomedicine, 2017: p. 305–315.

32. Grgacic, E.V. and D.A. Anderson, Virus-like particles: passport to immune recognition. Methods, 2006. 40(1): p. 60–65.

33. Choi, B., et al., Effective delivery of antigen–encapsulin nanoparticle fusions to dendritic cells leads to antigen-specific cytotoxic T cell activation and tumor rejection. ACS nano, 2016. 10(8): p. 7339–7350.

34. Szyszka, T.N., et al., High-Fidelity In Vitro Packaging of Diverse Synthetic Cargo into Encapsulin Protein Cages. Angewandte Chemie International Edition, 2025. 64(23): p. e202422459.s

35. Lohner, P., et al., Inside a shell—organometallic catalysis inside encapsulin nanoreactors. Angewandte Chemie, 2021. 133(44): p. 24028–24034.

36. Putri, R.M., et al., Structural characterization of native and modified encapsulins as nanoplatforms for in vitro catalysis and cellular uptake. ACS nano, 2017. 11(12): p. 12796–12804.

37. Shukla, S., Serum albumin ‘camouflage’of plant virus based nanoparticles prevents their antibody recognition and enhances pharmacokinetics. Biomaterials, 2016. 89: p. 89–97.

38. Pelaz, B., et al., Surface functionalization of nanoparticles with polyethylene glycol: effects on protein adsorption and cellular uptake. ACS nano, 2015. 9(7): p. 6996–7008.

39. Kaiser, C.R., et al., Biodistribution studies of protein cage nanoparticles demonstrate broad tissue distribution and rapid clearance in vivo. International journal of nanomedicine, 2007. 2(4): p. 715–733.

40. Singh, P., et al., Bio-distribution, toxicity and pathology of cowpea mosaic virus nanoparticles in vivo. Journal of controlled release, 2007. 120(1-2): p. 41–50.

41. Shukla, S., et al., Increased tumor homing and tissue penetration of the filamentous plant viral nanoparticle Potato virus X. Molecular pharmaceutics, 2013. 10(1): p. 33–42.

42. Bruckman, M.A., et al., Biodistribution, pharmacokinetics, and blood compatibility of native and PEGylated tobacco mosaic virus nano-rods and-spheres in mice. Virology, 2014. 449: p. 163–173.

43. Wang, J.-L., et al., The effect of surface poly (ethylene glycol) length on in vivo drug delivery behaviors of polymeric nanoparticles. Biomaterials, 2018. 182: p. 104–113.

44. Svergun, D.I., et al., Solution structure of poly (ethylene) glycol-conjugated hemoglobin revealed by small-angle X-ray scattering: implications for a new oxygen therapeutic. Biophysical journal, 2008. 94(1): p. 173–181.

45. Caballero, R.M., et al., Linear and multivalent PEGylation of the tobacco mosaic virus and the effects on its biological properties. Frontiers in Virology, 2023. 3: p. 1184095.

46. Pitek, A.S., et al., Serum albumin ‘camouflage’ of plant virus based nanoparticles prevents their antibody recognition and enhances pharmacokinetics. Biomaterials, 2016. 89: p. 89–97.

47. Li, M., et al., Brush conformation of polyethylene glycol determines the stealth effect of nanocarriers in the low protein adsorption regime. Nano letters, 2021. 21(4): p. 1591–1598.

48. Gómez-Barrera, S.N., et al., Surface engineering of the encapsulin nanocompartment of Myxococcus xanthus for cell-targeted protein delivery. Acs Omega, 2025. 10(7): p. 7142.

49. Bae, Y., et al., Engineering tunable dual functional protein cage nanoparticles using bacterial superglue. Biomacromolecules, 2018. 19(7): p. 2896–2904.

50. Choi, H., et al., Load and display: engineering encapsulin as a modular nanoplatform for protein-cargo encapsulation and protein-ligand decoration using split intein and SpyTag/SpyCatcher. Biomacromolecules, 2021. 22(7): p. 3028–3039.

51. Humphries, J., et al., Poly (2-oxazoline) and poly (2-oxazine)-based nanomedicines: Advancements, opportunities and challenges. Journal of Controlled Release, 2026: p. 114838.

52. Giessen, T.W., The structural diversity of encapsulin protein shells. Chembiochem, 2024. 25(24): p. e202400535.

53. Andreas, M.P. and T.W. Giessen, Large-scale computational discovery and analysis of virus-derived microbial nanocompartments. Nature Communications, 2021. 12(1): p. 4748.

