## Supplementary Information for "Modular PEGylation Confers Reduced Immune-Cell Uptake and Enhanced Pharmacokinetic Performance to Encapsulin Protein Nanocages"

**to Encapsulin Protein Nanocages**

Yi Wen Ng^1#^, Claire Rennie^1,2*#^, India Boyton^1,2^, Henrico Adrian^1^, Saluo Lai^3^,

Daryl Ariawan^4^, Peter Wich^3^, Ole Tietz^4^, Andrew Care^1,2,5*^

^1^School of Life Sciences, University of Technology Sydney, Gadigal Country, Sydney, NSW 2007, Australia

^2^Australian Institute for Microbiology and Infection, University of Technology Sydney, Gadigal Country, Sydney, NSW 2007, Australia

^3^School of Chemical Engineering, University of New South Wales, Sydney, NSW 2052, Australia

^4^Dementia Research Centre, Macquarie Medical School, Macquarie University, NSW 2109, Australia

^5^ARC Centre of Excellence in Synthetic Biology, Macquarie University, NSW 2109, Australia

#These authors contributed equally

Contents

**Supplementary Figure S1.** Purification of Am-S nanoscaffold

**Supplementary Figure S2.** Optimisation of Cy7 labelling conditions for Am-S

**Supplementary Figure S3A.** Chemical Characterization of Spytag Peptide

**Supplementary Figure S3B.** MALDI-TOF confirmation of S-PEG conjugation

**Supplementary Figure S4.** Optimisation of Am-S PEGylation

**Supplementary Figure S5.** Storage stability of Am-S^Cy7^ and Am-PEG^Cy7^

**Supplementary Figure S6.** Stability of Am-S^Cy7^ and Am-PEG^Cy7^ in serum-containing medium

**Supplementary Figure S7.** Serum-based calibration of Cy7-labelled encapsulins

**Supplementary Table S1.** PEG-conjugated protein nanoparticles investigated in cellular or *in vivo* models

**Supplementary Table S2.** Degree of labelling calculations for Am-S^Cy7^ at 1:0.3 Am-S to sulfo-Cy7-NHS ratio

**Supplementary Table S3.** Average polydispersity index (PDI) of Am-S^Cy7^ and Am-PEG^Cy7^ following freeze-thaw cycling and long-term storage

**Supplementary Table S4.** Blood sampling schedule for *in vivo* pharmacokinetic studies

**References**

**Abbreviations:**

| **Am-S** | SpyCatcher-displaying *Alkaliphilus metalliredigens* encapsulin |
| --- | --- |
| **Am-S^Cy7^** | Cy7-labelled Am-S |
| **Am-PEG^Cy7^** | PEGylated Cy7-labelled Am-S |
| **Am-PEG** | PEGylated Am-S |
| **Cy7** | Sulfo-Cyanine7 dye |
| **DLS** | Dynamic Light Scattering |
| **DMEM** | Dulbecco’s modified eagle medium |
| **FBS** | Fetal bovine serum |
| **IMAC** | Immobilised Metal Affinity Chromatography |
| **PEG** | Polyethylene glycol |
| **MALDI-TOF** | Matrix-Assisted Laser Desorption/Ionization Time-of-Flight |
| **SDS-PAGE** | Sodium dodecyl sulphate-polyacrylamide gel electrophoresis |
| **S-PEG** | PEGylated SpyTag |
| **SEC** | Size Exclusion Chromatography |

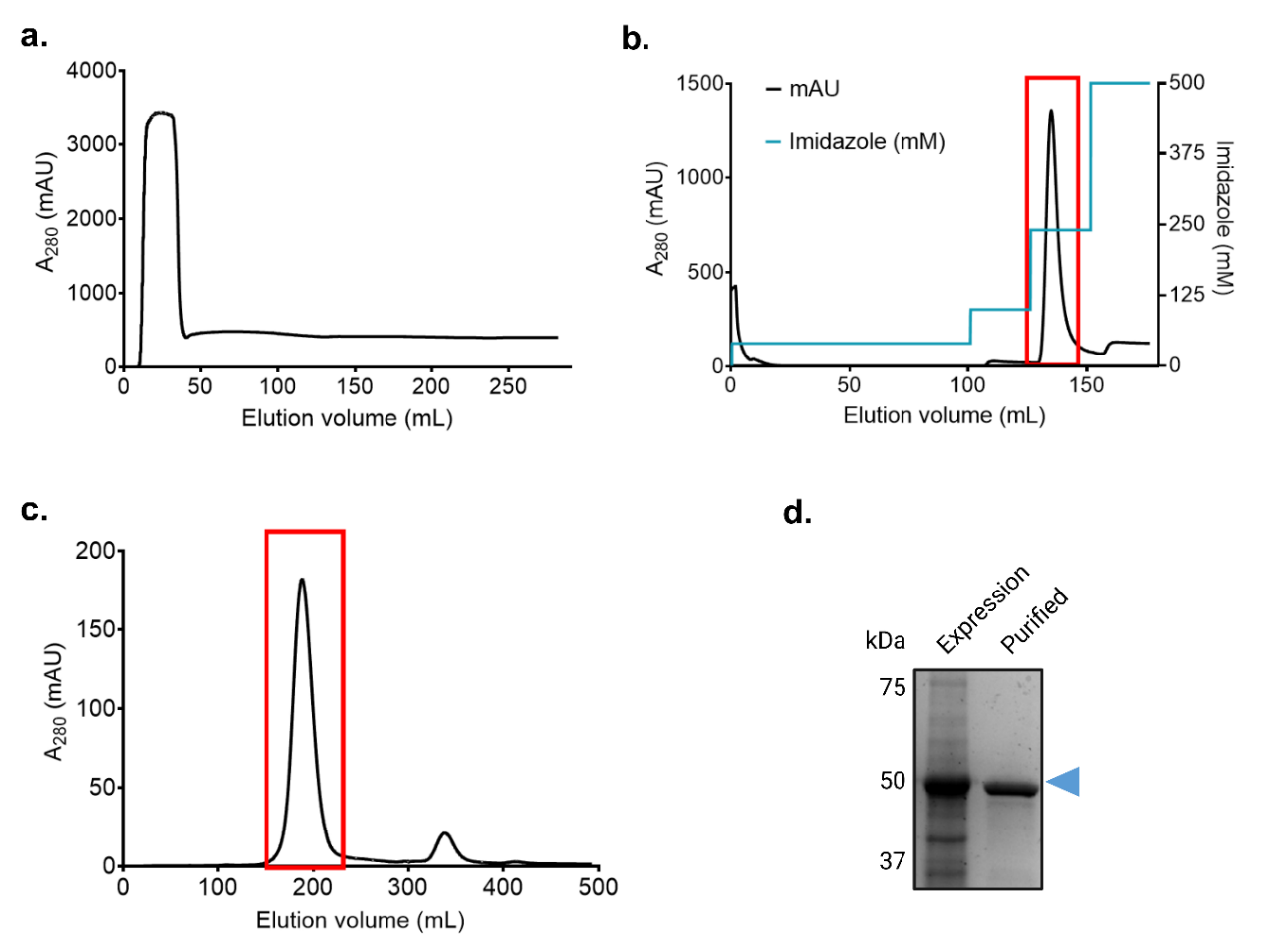

**Supplementary Figure S1. Purification of Am-S nanoscaffold.** Representative chromatogram of **(a)** on-column endotoxin removal during immobilised metal affinity chromatography (IMAC), **(b)** IMAC elution of Am-S, and **(c)** subsequent size-exclusion chromatography (SEC) purification. Am-S-containing peaks are highlighted in red squares. **(d)** Coomassie-stained SDS-PAGE showing Am-S expression and after final purification.

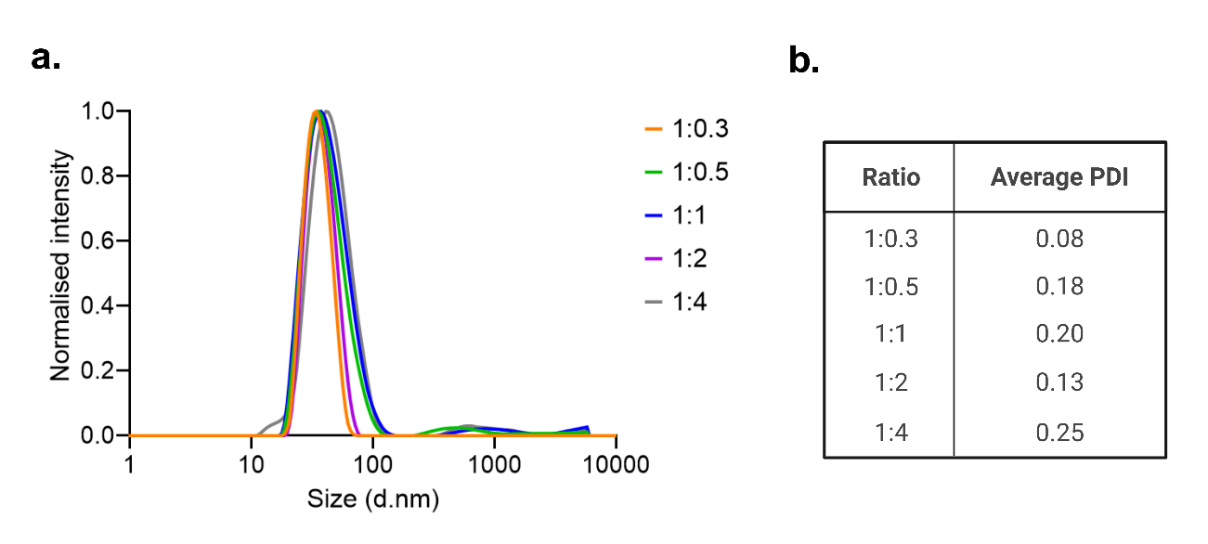

**Supplementary Figure S2. Optimisation of Cy7 labelling conditions for Am-S.** Dynamic light scattering (DLS) analysis of Am-S^Cy7^ prepared at varying Am-S:sulfo-Cy7-NHS conjugation ratios over a 6-day monitoring period. **(a)** DLS size distributions. **(b)** Average polydispersity index (PDI) values. Hydrodynamic diameter and PDI were monitored to assess the effect of Cy7 labelling density on colloidal stability. The 1:0.3 Am-S:sulfo-Cy7-NHS ratio remained colloidally stable throughout the study, whereas higher conjugation ratios exhibited increased PDI values and the appearance of aggregate populations.

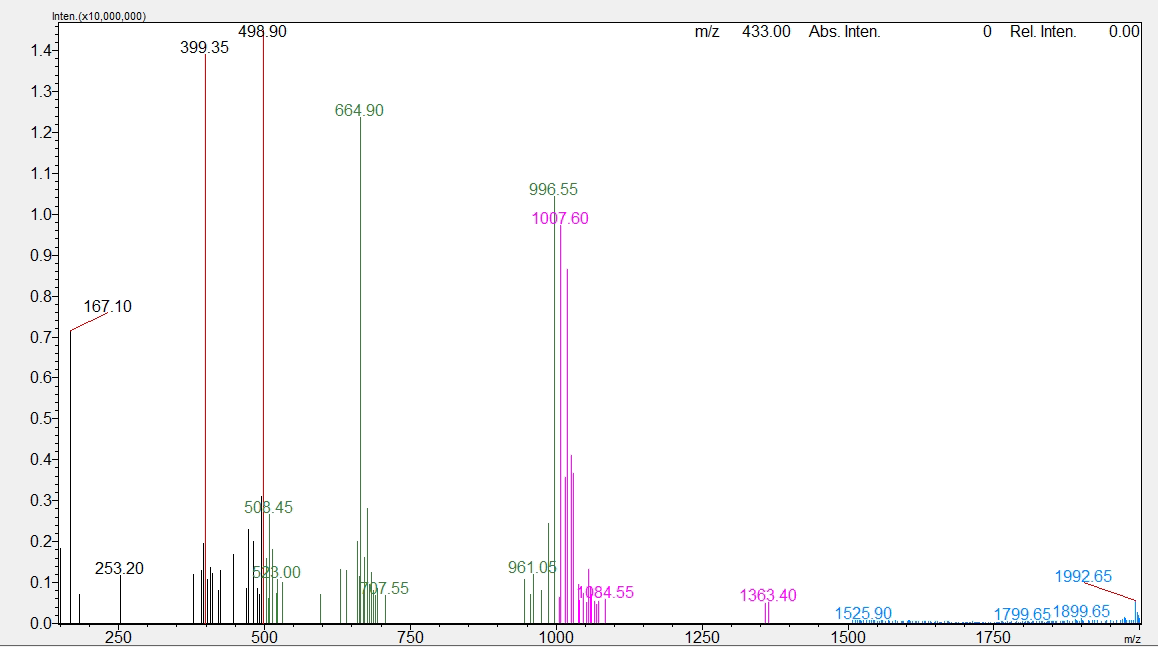

**Supplementary Figure S3A. Chemical Characterization of Spytag Peptide.** LCMS trace of Spytag peptide modified with a cysteine at the C-terminus

Sequence: AHIVMVDAYKPTKGSGSKC-NH_2_

Molecular Weight: 1991.36 g mol-1

LCMS [ESI+]:  1992.65 (m/z_1_; calculated: 1992.37), 996.55 (m/z_2_; calculated: 996.69), 664.90 (m/z_3_; calculated: 664.79), 498.90 (m/z_4_; calculated: 498.85), 399.35 (m/z_5_; calculated: 399.28)

HPLC retention time r_t_ = 13.8 min

Purity > 90 %

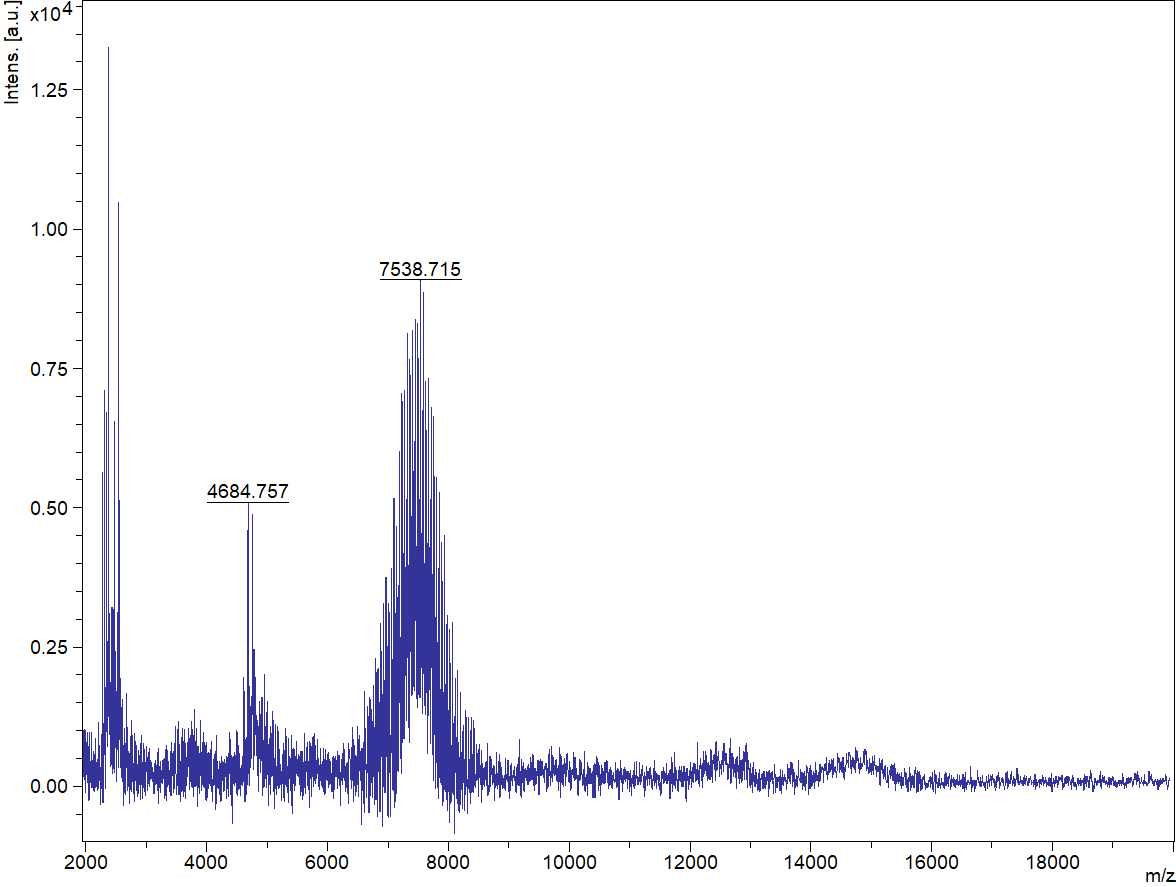

**Supplementary Figure S3B. MALDI-TOF confirmation of S-PEG conjugation.** SpyTag peptide (2.0 kDa) covalently conjugated to maleimide-functionalised PEG (4.5-5.0 kDa) through thiol–maleimide chemistry.

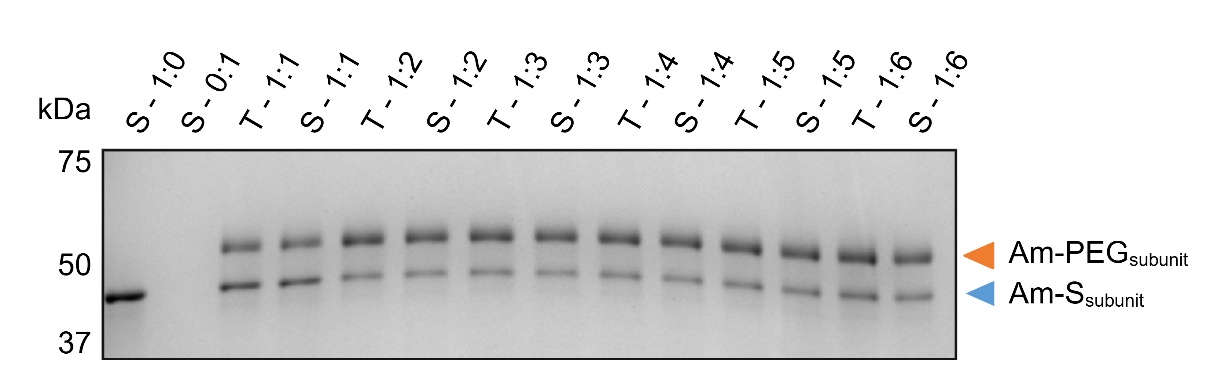

**Supplementary Figure S4. Optimisation of Am-S PEGylation.** Coomassie-stained SDS-PAGE of Am-S reacted with S-PEG at Am-S subunit:S-PEG molar ratios of 1:1 to 1:6. PEGylation was quantified by densitometry of PEGylated and unconjugated Am-S subunit bands. Subunit conversion increased with S-PEG ratio and plateaued at 1:3, reaching approximately 73%. S, soluble fraction; T, total fraction.

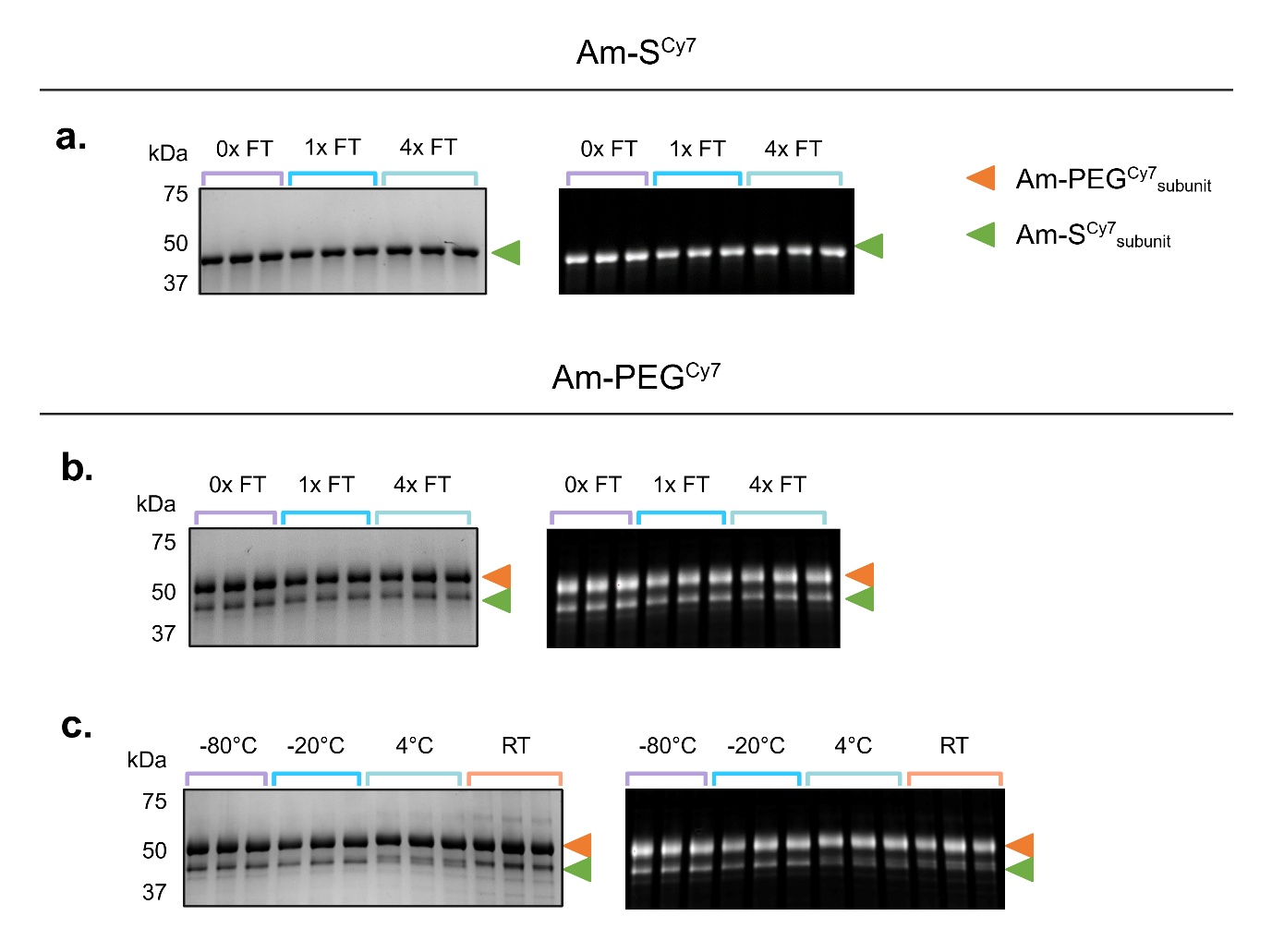

**Supplementary Figure S5. Storage stability of Am-S^Cy7^ and Am-PEG^Cy7^.** Coomassie-stained SDS-PAGE gels (left) and corresponding Cy7 fluorescence images (right) showing **(a)** Am-S^Cy7^ after 0, 1, and 4 freeze–thaw (FT) cycles, **(b)** Am-PEG^Cy7^ after 0, 1, and 4 FT cycles, and **(c)** Am-PEG^Cy7^ after 6 months of storage at −80 °C, −20 °C, 4 °C, or room temperature (RT). Gels were used to assess protein integrity and determine soluble protein recovery by densitometry.

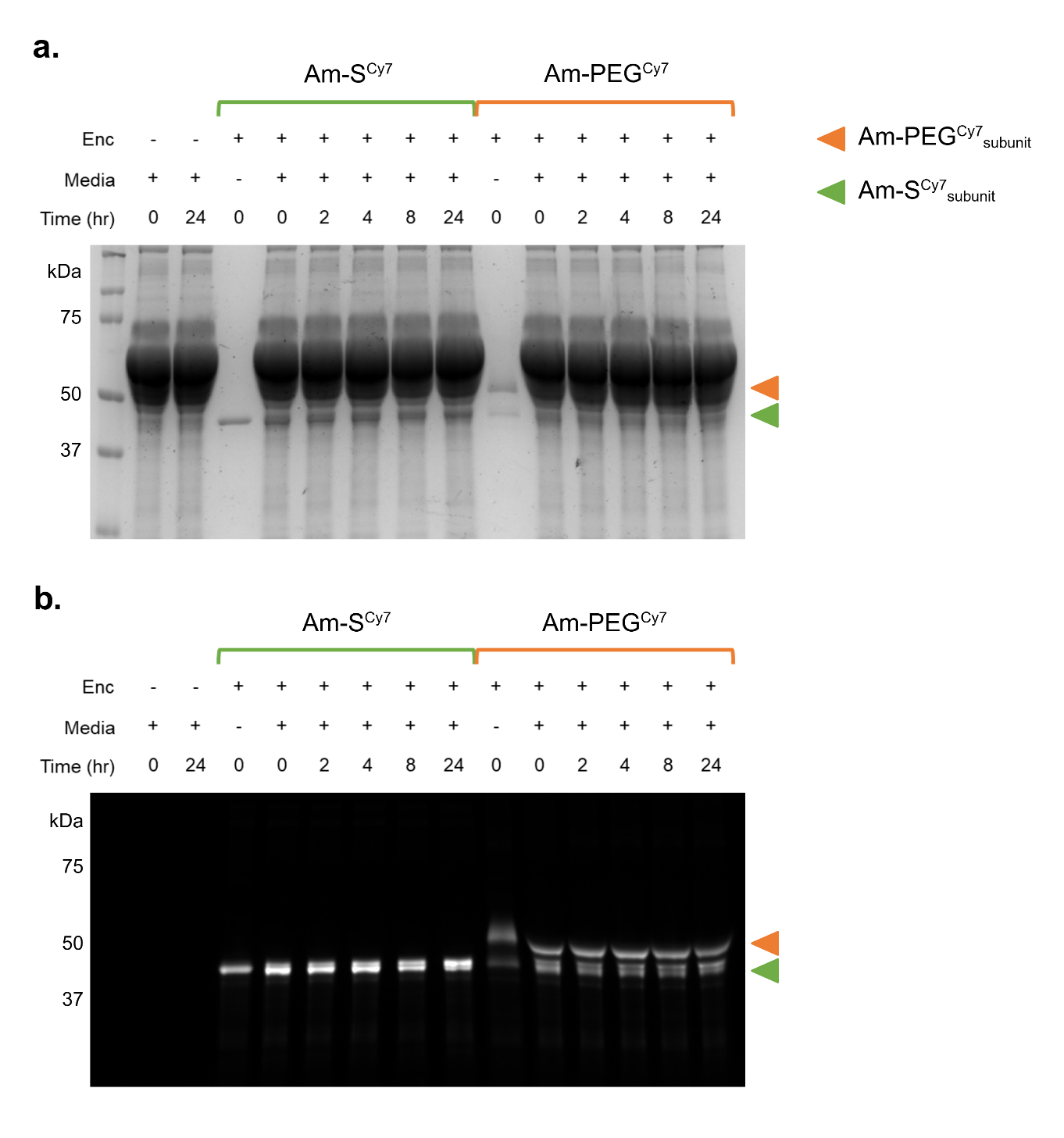

**Supplementary Figure S6. Stability of Am-S^Cy7^ and Am-PEG^Cy7^ in serum-containing medium. (a)** Coomassie-stained SDS-PAGE and **(b)** corresponding Cy7 fluorescence images of Am-S^Cy7^ and Am-PEG^Cy7^ following incubation in FBS-supplemented DMEM for 0, 2, 4, 8, and 24 h. Medium-only and encapsulin (Enc)-only controls are included.

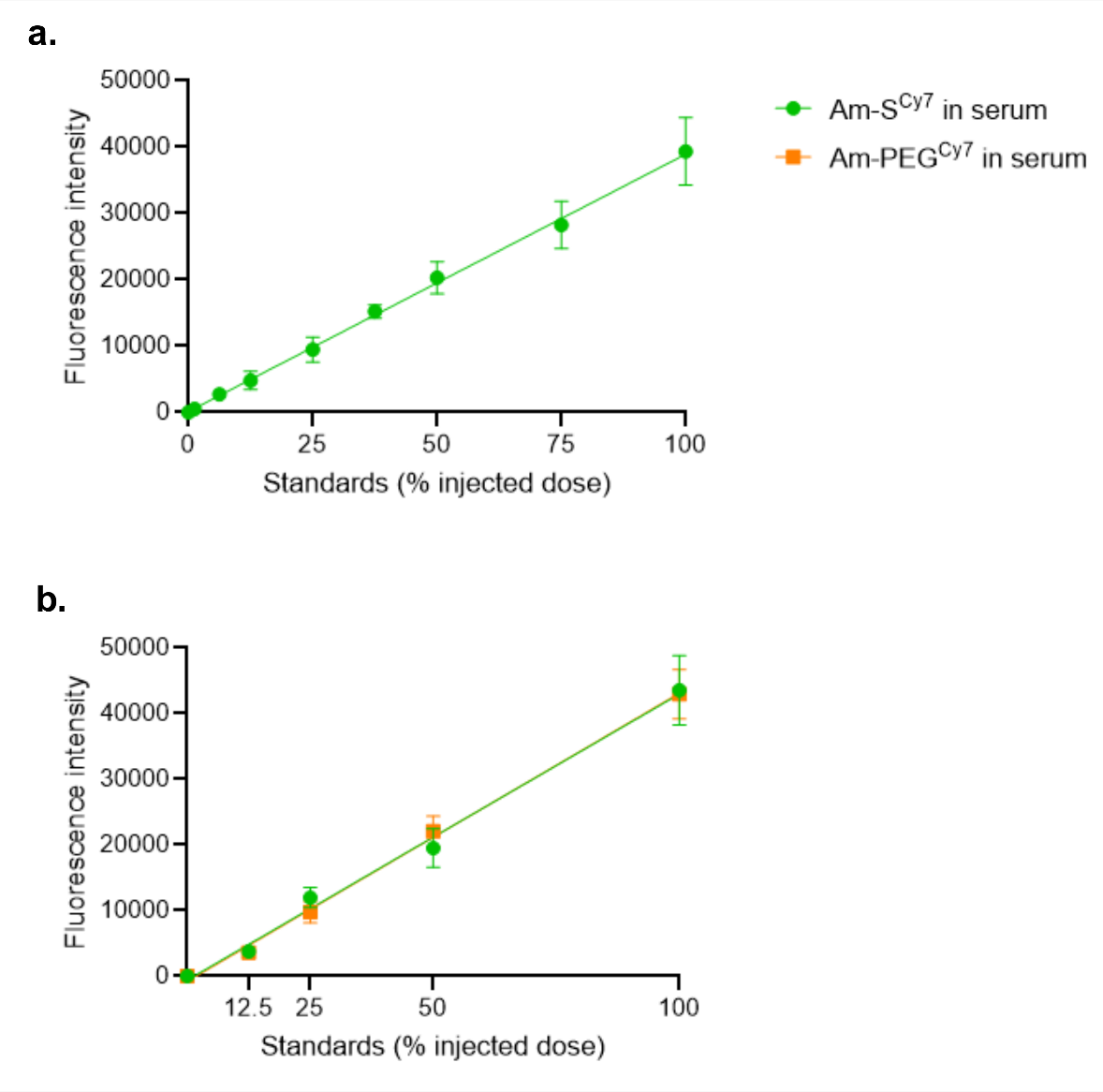

**Supplementary Figure S7. Serum-based calibration of Cy7-labelled encapsulins. (a)** Calibration curve generated by spiking naïve mouse serum with fluorescent Am-S^Cy7^ at concentrations corresponding to the indicated percentages of the injected dose. **(b)** Comparison of Am-S^Cy7^ and Am-PEG^Cy7^ fluorescence signals in naïve mouse serum, confirming comparable signal across the tested concentration range. Data are presented as mean ± SD (*n* = 3–5 for (a); *n* = 3 for (b)).

| **Supplementary Table S1. PEG-conjugated protein nanoparticles investigated in cellular or *in vivo* models** | | | | | | | | | |
| --- | --- | --- | --- | --- | --- | --- | --- | --- | --- |
| **PNP base** | **Conjugation strategy** | **Particle** | **Shape** | **Size** | **PEG length/conformation** | **pK T_1/2_ (mins)** | **Biodistribution** | **Other outcomes** | **Ref** |
| Cowpea Mosaic Virus (CPMV) | Lysine conjugation | CPMV | Sphere | 30 nm |  |  |  | Higher surface area grafting of PEG effectively prevented cell-particle interactions | [1] |
|  |  | CPMV-PEG1000 |  | 30 nm | MW 1000, mushroom |  |  |  |  |
|  |  | CPMV-PEG2000 |  | 30 nm | MW 20000, mushroom |  |  |  |  |
| Tobacco Mosaic Virus (TMV) | Lysine conjugation | TMV | Rod | 300 nm × 18 nm |  |  |  | PEG5000 shielded against anti-TMV antibodies. Antibody evasion decreased for multivalent PEG5000 | [2] |
|  |  | 2L |  |  | MW 2000, linear |  |  |  |  |
|  |  | 2B |  |  | MW 2000, bivalent |  |  |  |  |
|  |  | 2R |  |  | MW 2000, 4arm |  |  |  |  |
|  |  | 5L |  |  | MW 5000, linear |  |  |  |  |
|  |  | 5B |  |  | MW 5000, bivalent |  |  |  |  |
|  |  | 5R |  |  | MW 5000, 4arm |  |  |  |  |
| Tobacco Mosaic Virus (TMV) | Side chain cycloaddition | Cy5-TMV | Rod | 300 nm × 18 nm |  | Fast - 3.5, slow - 94.9 mins | 4 h - liver, spleen  24 h - liver, spleen  96 h - stomach |  | [3] |
|  |  | PEG-Cy5-TMV | Rod | 300 nm × 18 nm | MW 2000 | Fast - 6.3, slow - 44.4 mins | 4 h - liver, spleen  24 h - liver, spleen  96 h - spleen |  |  |
| Tobacco Mosaic Virus (TMV) | Lysine conjugation | TMV-PEG_24_ | Rod | 300 nm × 18 nm | 1394.55, mushroom | 10 mins |  | PEGylation decreased interaction with macrophages *in vitro* | [4] |
|  |  | TMV-PEG_105_ |  |  | 5000, mushroom |  |  |  |  |
| Potato Virus X (PVX) | Lysine conjugation | A-PVX | Filamentous |  |  | 19 mins | Whole body until 8 h, then liver and spleen until 8 h, then cleared | P5B and P20 showed reduced tissue accumulation, consistent with extended circulation times, as expected from PEGylation | [5] |
|  |  | A-PVX-P5L | Filamentous |  | MW 5000, linear, mushroom | Fast – 11, slow – 409 mins |  |  |  |
|  |  | A-PVX-P5B | Rod |  | MW 5000, branched, mushroom | Fast – 14, slow – 1142 mins | Accumulation primarily in liver and spleen only up to 8 h |  |  |
|  |  | A-PVX-P20 | Rod |  | MW 20000, brush | Fast – 27, slow – 231 mins |  |  |  |
| Physalis Mottle Virus (PhMV) | Cysteine conjugation | PhMV | Sphere | 30.7 nm |  |  |  |  | [6] |
|  |  | Gd-Cy5.5-PhMV-mPEG | Sphere | ~33.6 nm | MW 2000 | Fast – 133, slow – 2616 mins | Uptake to liver, spleen and tumour significantly less than a targeted particle |  |  |
| Cowpea Mosaic Virus (CPMV) and Potato Virus X (PVX) | Lysine conjugation | A647-CPMV-PEG | Sphere | 30 nm | MW 5000 | 20.8 mins | Accumulated primarily in the liver at 24 h (71%), spleen (13%) and tumour (9%) |  | [7] |
|  |  | A647-PVX-PEG | Filamentous | 515 nm x 13 nm | MW 5000 | 12.5 mins | Primarily accumulated in the liver but to a lesser degree (55%). More accumulated in the spleen (23%) and tumour (15%) |  |  |
| Major capsid protein of simian virus 40 (SV40) | Lysine conjugation | Ag2S@PNC_SV40_ | Sphere | 24 ± 1.76 nm |  | <5 | High accumulation in the spleen 12h post-injection |  | [8] |
|  |  | Ag2S@PNC_SV40_-PEG750 |  |  | MW 750 | 283 mins | Minimal accumulation in tissues consistent with the properties of PEG |  |  |
|  |  | Ag2S@PNC_SV40_-PEG5K |  |  | MW 5000 | 476 mins |  |  |  |
| Ferritin | Lysine conjugation | FTn | Sphere | 10.5 nm |  | 25.2 mins |  |  | [9] |
|  |  | PEG-FTn 75% |  | 13.3 nm | MW 2000 | 73.8 mins | Accumulated in the tumours over time |  |  |

**Supplementary Table S2. Degree of labelling calculations for Am-S^Cy7^ at 1:0.3 Am-S to sulfo-Cy7-NHS ratio.**

| A280 | 0.702 |
| --- | --- |
| A750 | 0.667 |
| Correction factor | 0.04 |
| Dilution factor | 1 |
| Ɛ of protein | 48360 L·mol⁻¹·cm⁻¹ |
| Ɛ of dye | 240600 L·mol⁻¹·cm⁻¹ |
| Protein concentration | 1.40 x 10^-5^ M |
| Dye:protein molar ratio | 0.199 |
| DOL per nanocage (60 subunits) | ~12 Cy7/cage |

**Supplementary Table S3. Average polydispersity index (PDI) of Am-S^Cy7^ and Am-PEG^Cy7^ following freeze-thaw cycling and long-term storage.**

| **Stability Assessment** | **Construct** | **Condition** | **Average PDI** |
| --- | --- | --- | --- |
| Freeze-thaw | Am-S^Cy7^ | 0x FT | 0.05 |
|  |  | 1x FT | 0.13 |
|  |  | 4x FT | 0.17 |
|  | Am-PEG^Cy7^ | 0x FT | 0.06 |
|  |  | 1x FT | 0.07 |
|  |  | 4x FT | 0.08 |
| Long-term storage | Am-PEG^Cy7^ | -80°C | 0.07 |
|  |  | -20°C | 0.16 |
|  |  | 4°C | 0.08 |
|  |  | RT | 0.10 |

**Supplementary Table S4. Blood sampling schedule for *in vivo* pharmacokinetic studies.** Non-terminal blood samples were collected *via* submandibular bleeds at early timepoints (1, 3, and/or 6 hours), with a maximum of two non-terminal bleeds per mouse. Terminal blood collection *via* cardiac puncture was performed at 12- or 24-hours post-injection. Mouse 1 was found deceased shortly before the scheduled 3-hour timepoint; as the 3-hour blood sample had been collected, this data point was retained. The cause of death was undetermined. Mice 3 and 11 experienced missed injections and were excluded from all pharmacokinetic analyses. Mice 8, 9, 28, 29, and 30 received no treatment and were designated as untreated (naïve) controls.

○ = submandibular bleed.	● = terminal cardiac puncture.

| Mouse # | Treatment | Timepoint (hour) | | | | |
| --- | --- | --- | --- | --- | --- | --- |
|  |  | 1 | 3 | 6 | 12 | 24 |
| 1 | Am-S^Cy7^ | ○ | ● |  |  |  |
| 2 | Am-S^Cy7^ | ○ |  | ○ | ● |  |
| 3 | Missed |  |  |  |  | ● |
| 4 | Am-S^Cy7^ | ○ |  |  | ○ | ● |
| 5 | Am-S^Cy7^ | ○ |  | ○ | ● |  |
| 6 | Am-S^Cy7^ |  | ○ | ○ |  | ● |
| 7 | Am-S^Cy7^ |  |  | ○ |  | ● |
| 8 | Control |  |  |  | ● |  |
| 9 | Control |  |  |  | ● |  |
| 10 | Am-S^Cy7^ | ○ | ○ |  | ● |  |
| 11 | Missed |  |  |  |  | ● |
| 12 | Am-PEG^Cy7^ |  |  | ○ |  | ● |
| 13 | Am-S^Cy7^ |  | ○ | ○ |  | ● |
| 14 | Am-S^Cy7^ |  | ○ |  | ○ | ● |
| 15 | Am-S^Cy7^ |  |  | ○ |  | ● |
| 16 | Am-S^Cy7^ | ○ |  |  | ○ | ● |
| 17 | Am-S^Cy7^ |  | ○ | ○ | ● |  |
| 18 | Am-PEG^Cy7^ | ○ | ○ |  | ● |  |
| 19 | Am-PEG^Cy7^ | ○ |  | ○ | ● |  |
| 20 | Am-PEG^Cy7^ | ○ | ○ |  | ● |  |
| 21 | Am-PEG^Cy7^ | ○ |  |  | ○ | ● |
| 22 | Am-PEG^Cy7^ | ○ |  | ○ | ● |  |
| 23 | Am-PEG^Cy7^ |  | ○ | ○ |  | ● |
| 24 | Am-PEG^Cy7^ |  | ○ | ○ |  | ● |
| 25 | Am-PEG^Cy7^ |  | ○ |  |  | ● |
| 26 | Am-PEG^Cy7^ |  | ○ |  |  | ● |
| 27 | Am-PEG^Cy7^ |  |  | ○ |  | ● |
| 28 | Control |  |  |  | ● |  |
| 29 | Control |  |  |  | ● |  |
| 30 | Control |  |  |  | ● |  |

**References**

1.	Steinmetz, N.F. and M. Manchester, *PEGylated viral nanoparticles for biomedicine: the impact of PEG chain length on VNP cell interactions in vitro and ex vivo.* Biomacromolecules, 2009. **10**(4): p. 784-792.

2.	Caballero, R.M., et al., *Linear and multivalent PEGylation of the tobacco mosaic virus and the effects on its biological properties.* Frontiers in Virology, 2023. **3**: p. 1184095.

3.	Bruckman, M.A., et al., *Biodistribution, pharmacokinetics, and blood compatibility of native and PEGylated tobacco mosaic virus nano-rods and-spheres in mice.* Virology, 2014. **449**: p. 163-173.

4.	Pitek, A.S., et al., *Serum albumin ‘camouflage’ of plant virus based nanoparticles prevents their antibody recognition and enhances pharmacokinetics.* Biomaterials, 2016. **89**: p. 89-97.

5.	Lee, K.L., et al., *Stealth filaments: Polymer chain length and conformation affect the in vivo fate of PEGylated potato virus X.* Acta biomaterialia, 2015. **19**: p. 166-179.

6.	Hu, H., et al., *Physalis mottle virus-like nanoparticles for targeted cancer imaging.* ACS applied materials & interfaces, 2019. **11**(20): p. 18213-18223.

7.	Shukla, S., et al., *Increased tumor homing and tissue penetration of the filamentous plant viral nanoparticle Potato virus X.* Molecular pharmaceutics, 2013. **10**(1): p. 33-42.

8.	Li, C., et al., *Real-time monitoring surface chemistry-dependent in vivo behaviors of protein nanocages via encapsulating an NIR-II Ag2S quantum dot.* Acs Nano, 2015. **9**(12): p. 12255-12263.

9.	Huang, X., et al., *Hypoxia-tropic protein nanocages for modulation of tumor-and chemotherapy-associated hypoxia.* Acs Nano, 2018. **13**(1): p. 236-247.
